# Dorsal anterior cingulate-midbrain ensemble as a reinforcement meta-learner

**DOI:** 10.1101/130195

**Authors:** Massimo Silvetti, Eliana Vassena, Elger Abrahamse, Tom Verguts

**Affiliations:** Ghent University, Department of Experimental Psychology; Donders Institute for Brain, Cognition and Behaviour

**Keywords:** ACC, VTA, LC, reinforcement learning, meta-learning, Kalman filter, effort control, higher-order conditioning, decision-making

## Abstract

The dorsal anterior cingulate cortex (dACC) is central in higher-order cognition and behavioural flexibility. The computational nature of this region, however, has remained elusive. Here we propose a new model – the Reinforcement Meta Learner (RML) – based on the bidirectional anatomical connections of the ACC with midbrain catecholamine nuclei (VTA and LC). In this circuit, dACC learns which actions are valuable and acts accordingly. Crucially, this mechanism is optimized by recurrent connectivity with the midbrain: Midbrain catecholamines provide modulatory signals to dACC, controlling its internal parameters (e.g. learning rate), while these parameter modulations are in turn optimized by dACC afferents to the midbrain. This closed-loop system generates emergent (i.e., homunculus-free) control and supports learning to solve hierarchical decision problems without having an intrinsic hierarchical structure itself. Further, it can be combined with other cortical modules to optimize the processing of these modules. We outline how the RML solves the current theoretical stalemate on dACC by assimilating various previous proposals on ACC functioning, and how it captures critical empirical findings from an unprecedented range of domains (stability/plasticity balance, effort processing, working memory, and higher-order classical and instrumental conditioning).

## Introduction

Adapting behavior to uncertain and changing environments is the foundation of intelligence. Important theoretical progress was made by considering this behavioural adaptation as a problem of decision-making (Frank et al., 2004; Rushworth and Behrens, 2008). The dorsal anterior cingulate cortex (dACC) has been proposed as a multifunctional hub with a pivotal role in decision-making (Rushworth and Behrens, 2008). Reinforcement Learning (RL) neural models showed that many signals recorded in the dACC (e.g., error, error likelihood, response conflict) can be explained in terms of value computation for optimal decision-making (Silvetti et al., 2014). In this framework, the dACC is a multi-domain estimator of stimulus and action values that maximizes long-term reward. Yet, it is increasingly clear that value computation by itself cannot fully account for dACC functioning. Besides computing environment- and behavior-related parameters (e.g., outcome prediction, etc.), dACC is also involved in adaptive control over the organism’s internal cognitive parameters (e.g., learning rate). Empirical evidence indeed shows that dACC controls effort exertion (Shenhav et al., 2013; Vassena et al., 2014; Verguts et al., 2015). Moreover, a Bayesian perspective proposes that the dACC controls learning rate to optimize behavioural adaptation (Behrens et al., 2007; Kolling et al., 2016). Despite the lively ongoing debate on these computational functions and empirical effects measured in ACC no theoretical convergence has been reached so far (Ebitz and Hayden, 2016), and no concrete computational model has been developed to reconcile (or allow competition between) such different theoretical positions (see Vassena et al., 2017 for a review).

To fill this lacuna, we here propose a novel RL model coined the Reinforcement Meta Learner (RML). To achieve a comprehensive understanding of dACC functioning, we argue that it is critical to place dACC in its larger functional network. The RML model is the first to theoretically build on the demonstrated bidirectional anatomical connections between the ACC and the midbrain catecholamine nuclei (Devinsky et al., 1995; Margulies et al., 2007), the ventral tegmental area (VTA), and the locus coeruleus (LC). Like in earlier RL models, the dACC in RML computes the values of specific stimuli and actions to achieve adaptive behavior. However – and unlike earlier models – dACC internal parameters are dynamically set by midbrain catecholamine nuclei. Crucially, VTA and LC outputs are in turn determined by dACC, implementing a recurrent *meta-learning* architecture (Figure 1a).

**Figure 1.**
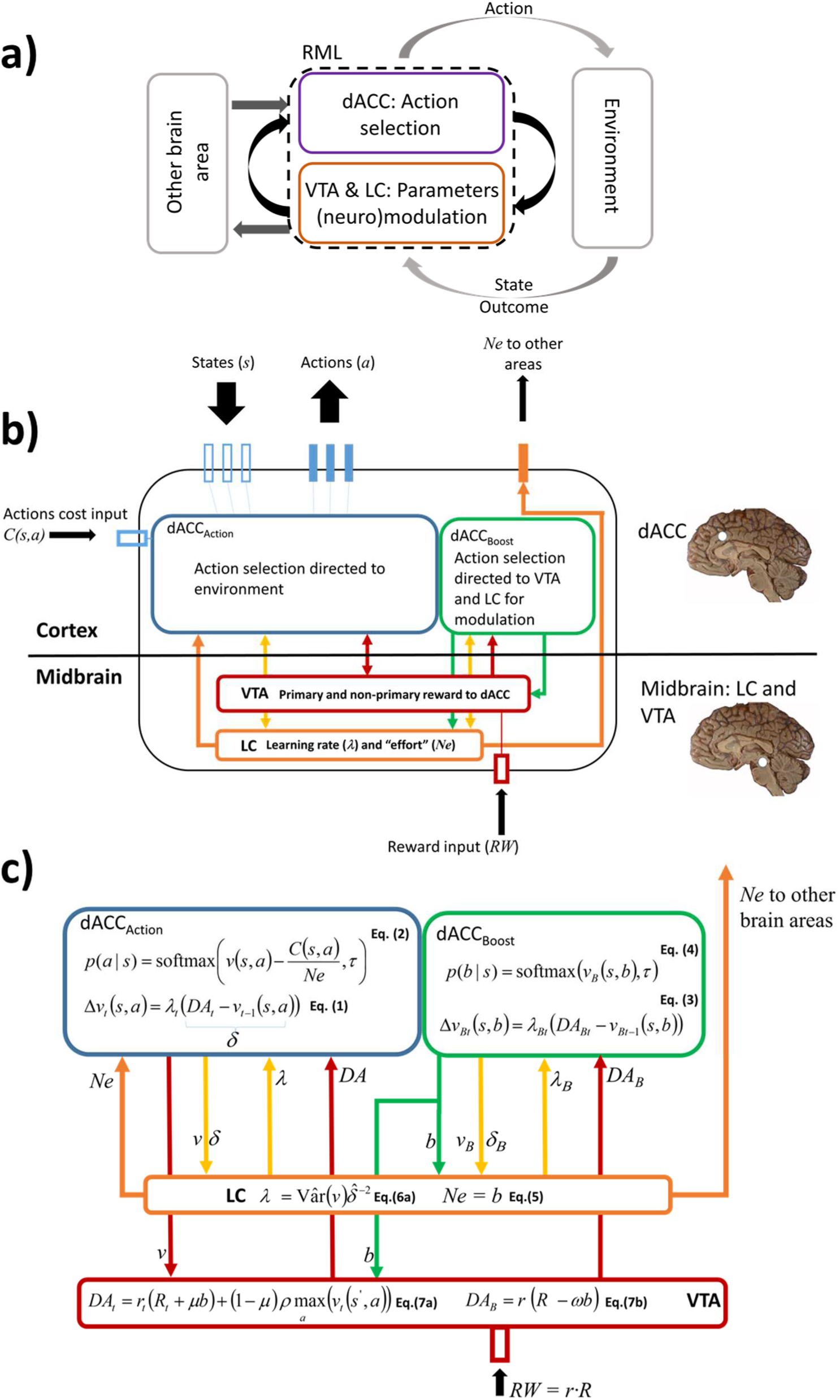
a) Model conceptual overview. At the conceptual level, the RML dynamics can be summarized in three recurrent circuits. First, a main internal loop (black arrows) indicating recurrent interaction between action selection processes and parameter control processes, which together determine the emergence of meta-learning. Second, an external loop (light grey arrows), indicating the interactions between the RML and environment. Third, a secondary (optional) internal loop (dark grey), determining the interactions between the RML and other cognitive modules, receiving catecholamines input from the RML and providing back a signal influencing the decision-making processes in the RML itself. **b) Model overview with anatomical-functional analogy.** The RML-environment interaction happens through nine channels of information exchange (black arrows) (input = empty bars; output = filled bars). The input channels consist of one channel encoding action costs (*C*), three encoding environmental states (*s*) and one encoding primary rewards (*RW*). The output consists of three channels coding each for one action (*a*), plus one channel conveying LC signals to other brain areas (*Ne*). The entire model is composed of four reciprocally connected modules (each a different color). The upper modules (blue and green) simulate the dACC, while the lower modules (red and orange) simulate the midbrain catecholamine nuclei (VTA and LC). dACC_Action_ selects actions directed toward the environment and learns through first and higher-order conditioning, while dACC_Boost_ modulates catecholamine nuclei output. The VTA module provides DA training signals to both dACC modules, while the LC controls learning rate (yellow bidirectional arrow) in both dACC modules, and effort exertion (promoting effortful actions) in the dACC_Action_ module (orange arrow), influencing their decisions. Finally, the LC signal controlling effort in the dACC_Action_ is directed also toward other brain areas for neuro-modulation. **c) Model overview with equations.** The equations are reported in their discrete form. Communication between modules is represented by arrows, with corresponding variables near each arrow. Variables *δ* and *δ*_*B*_ represent the prediction errors from respectively equations 1 and 3. For the complete description of equations and symbols, refer to the Model Description: mathematical formalization section below.

The RML innovates ACC theory in two major ways. First, interaction with both the environment and the midbrain, allows the dACC to modulate autonomously three critical variables: i.e. reward, effort, and learning rate. This closed-loop system solves hierarchical decision problems (i.e. about the context on where a task is executed) without having an intrinsic hierarchical structure itself. For example, selection of an optimal learning rate is conditional on the estimation of volatility, a hidden variable that determines the statistical structure of task context (Silvetti et al., 2013). As another example, the model decides how much effort to put in a task, as a function of the overall favorability (in terms of reward) of the context.

Moreover, in the RML these three meta-learning processes (reward, effort and learning rate) interact with one another, optimizing cognitive flexibility. For example, learning rate changes the plasticity of effort modulation, and effort modulation changes the interaction with the environment (modifying the preference toward effortful options), which in turn influences learning rate.

Second, the dACC-midbrain ensemble can be considered a general provider of control signals to other brain areas (via Ne output). The RML model can be plugged into other models to modulate them with this functionality. As such, the RML fully exploits both *modularity* (between models) and distributed computing (dACC-midbrain loop), accounting for how cognitive control flexibly adapts across different tasks. To the best of our knowledge, the RML is the first model that can improve the performance of different, independently designed and published models.

Thanks to these features, the RML captures critical empirical findings from an unprecedented range of domains, namely research on the stability/plasticity balance, on effort processing, on working memory, and on higher-order classical and instrumental conditioning. Critically, besides the above outlined notion that the RML dynamically estimates three of its parameters, for the remaining parameters a single set is used across all domains and simulations – thus without experimenter-based tuning. In the next section, we take a closer look at the RML model.

## Model description: overview

Here we describe the neuro-functional architecture of the RML model. We present the different modules, their neuroanatomical correlates, and the proposed connectivity patterns, followed by the computational mechanisms implemented in each module. Importantly, we carried out two different implementations of the model: a discrete version, modeling trial by trial changes, and a continuous version, modeling brain activity with ten milliseconds steps. The discrete-time description was chosen to facilitate understanding of the model’s conceptual structure; yet, a continuous-time (dynamical) implementation was necessary to study intra-trial dynamics, like DA shifting from US to CS or working memory processing. For simplicity, we here describe the discrete version and refer the reader to the Supplementary Methods for description of the continuous version. Although we ran all the simulations not requiring intra-trial dynamics also with the discrete model, leading to equivalent results (see Supplementary Results: main results replication with discrete model), the simulations described in the Results section are from the dynamical version of RML.

The RML approximates optimal decision-making (to maximize long-term reward) based on action-outcome comparisons. Model dynamics can by summarized by Figure 1a, where three loops are described: an inner loop (between action-selection and parameter control processes, black arrows), and an external loop, between RML and environment (light grey arrows). The external loop directly influences both parameter control module (through primary rewards) and action selection module (through state transitions), for this reason it can be considered a second communication channel between action selection and parameter control modules. As described more in detail below, both loops contribute to the emergence of meta-learning dynamics. Finally, a third loop simulates how the RML can exert control over other brain areas, via catecholaminergic signals, and how these areas can modulate back the decision-making processes in the RML. The latter loop is not intrinsically necessary for the RML functioning, although it shows how the dACC-midbrain system can work as a “server” providing control signals to optimize behavioural performance.

An overview closer to neurophysiology (Figure 1b) shows that action-selection and parameter control belong respectively to cortical and midbrain structures: The dACC (blue and green modules in Figure 1b) performs action-outcome comparison and action selection, while it is augmented by meta-learning (LC and VTA; orange and red modules in Figure 1b). We designed the model such that the communication with the external environment is based on 9 channels. There are six channels representing environmental states and RML actions (3 states and 3 actions). The first two actions are aimed at changing the environmental state (e.g. turning right or left), while the 3^rd^ action means “Stay”, i.e. refusing to engage in the task. There are two other input channels, one dedicated to primary reward from environment and the other to signal indicating costs of motor actions. Finally, there is one output channel conveying norepinephrine (*Ne*) signals to other brain areas (cognitive control signal).

In Figure 1c we link the model equations described below (“Model description: mathematical formalization”) with a graphical representation. As we describe in detail in the following section, the RML autonomously modulates three different parameters providing a near optimal solution for the following meta-learning problems. First, modulation of *learning rate* (*λ*) ensures that knowledge is updated only when there are relevant environmental changes, protecting it from non-informative random fluctuations. This addresses the classical stability-plasticity trade-off (Grossberg, 1980). Second, modulation of effort exertion (by control over *N*e) allows optimization when the benefit-cost balance changes over time and/or it is particularly challenging to estimate (e.g. when it is necessary to exert high effort to get an uncertain higher reward). Third, dynamic modulation of *reward* signals (by means of control over *DA*) is the foundation for emancipating learning from primary rewards (see Equation 7 and Equation s5), allowing the RML to learn complex tasks without the immediate availability of primary rewards (higher order conditioning).

Moreover, adaptive *DA* signal implies modulating the (perceived) value of state/action couples (*DA* represents outcome in Equation 1), eventually self-motivating the RML in choosing one specific action even if it implies a high execution cost (*C*, Equation 2), and thus influencing the RML behaviour also in challenging cost-benefit problems. Meta-learning processes involving those three variables (learning rate, effort, reward) interact with each other, based on the dynamics described below, and thus forming an integrated system where flexibility emerges from its recurrent dynamics.

The RML is scalable by design, i.e. there is no theoretical limit to the number of state/action channels, and neither the number of parameters nor their values changes as a function of task type/complexity. As specified below, we used a single set of parameters across all simulations both for the discrete model (Table 1) and for the dynamic model (see Table s1).

**Table 1.** Parameters list and values for the discrete model. We ran all the simulations using one set of parameter values (for both the discrete and dynamical model).

| Parameter | Value | Meaning | Equation |
| --- | --- | --- | --- |
| $\rho$ | 0.2 | TD-learning signal decay | 7a |
| $\mu$ | 0.3 | DA dynamics | 7a |
| $\tau$ | 0.6 | Softmax temperature | 2 |
| $\alpha$ | 0.3 | Kalman filtering meta-parameter | 6c-d |
| $\beta$ | 0.2 | Learning rate lower bound | 6 |
| $\omega$ | 0.15 | Boosting cost | 7b |

## Model description: mathematical formalization

### dACC_Action_

The dACC_Action_ module consists of an Actor-Critic system augmented with meta-learning (Figure 1c, blue box). The Critic is a performance evaluator and computes reward expectation and PE for either primary or non-primary rewards (higher-order conditioning), learning to associate stimuli and actions to environmental outcomes. The Actor selects motor actions (based on Critic expectation) to maximize long-term reward.

The central equation in this module governs Critic state/action value updates:

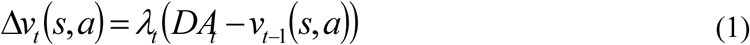

where *v*(*s*,*a*) indicates the value (outcome prediction) of a specific action *a* given a state *s*. Equation 1 ensures that *v* comes to resemble the environmental outcome encoded by dopaminergic signal (*DA*), which is generated by the VTA module (Figure 1; Equation 7). It entails that the update of *v* at trial *t* is based on the difference between prediction (*v*) and outcome (*DA*), which defines the concept of PE. The latter is weighted by learning rate *λ*, making the update more (high *λ*) or less (low *λ*) dependent on recent events. We propose that *λ* itself is modulated by the LC based on *v* and PE signals from the dACC_Action_ (Equation 6a).

The *DA* signal, afferent from the VTA, is either linked to primary or non-primary reward (higher-order conditioning) and is modulated by the dACC_Boost_ module via parameter *b* (Equation 7a).

Action *a* is selected by the Actor subsystem, which implements action selection (by softmax selection function, with temperature *τ*) based on state/action values discounted by state/action costs *C*:

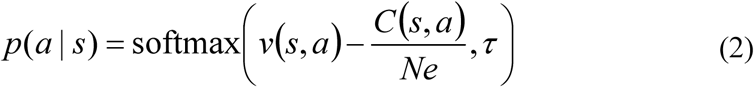

Function *C* assigns a cost to each state/action couple, for example energy depletion consequent to climbing an obstacle. *C* is modulated by norepinephrine afferents from LC (*Ne*), which is itself controlled by the dACC_Boost_ module, via parameter *b. Ne* levels discount *C*, lowering the perceived costs and energizing behaviour. Here we remind the reader that the RML can choose not to engage the task (“Stay”); this option has *C* = 0. In this way, a high level of *Ne* energizes behaviour, promoting both high cost actions and reducing the probability that the RML chooses to “Stay”.

The dynamical form of these equations is described in the *dACC*_*Action*_*-VTA system* paragraph in Supplementary Methods.

### dACC_Boost_

The dACC_Boost_ module is an Actor-Critic system that learns only from primary rewards (Figure 1c, green box). This module controls the parameters for cost and reward signals in equations 1-2 (dACC_Action_), via modulation of VTA and LC activity (boosting catecholamines). In other words, whereas the dACC_Action_ decides on actions toward the external environment, the dACC_Boost_ decides on actions toward the internal environment: It modulates midbrain nuclei (VTA and LC), given a specific environmental state. This is implemented by selecting the modulatory signal *b* (*boost signal*), by RL-based decision-making. In our model, *b* is a discrete signal that can assume ten different values (integers 1-10), each corresponding to one action selectable by the dACC_Boost_. The Critic submodule inside the dACC_Boost_ updates the boost values *v*_*B*_(s,b), via the equation:

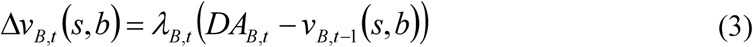

Equation 3 represents the value update of boosting level *b* in the environmental state *s*. The dACC_Boost_ module receives dopaminergic outcome signals (*DA*_*B*_) from the VTA module. As described in Equation 7b, *DA*_*B*_ represent the reward signal discounted by the cost of boosting catecholamines (Kool et al., 2010; Kool and Botvinick, 2013; Shenhav et al., 2013). Also in Equation 3 there is a dynamic learning rate (*λ*_*B*_), estimated by Equation 6a in the LC. The Actor submodule selects boosting actions to maximize long-term reward:

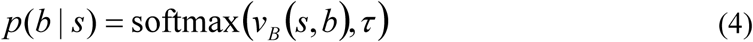

Referring to Equation 1, the dACC_Boost_ modulates the reward signal by changing the *DA* signal coded in VTA (Equation 7a). Furthermore, dACC_Boost_ also modulates the cost signal by changing parameter *Ne* (via LC module, Equation 5) in the function representing action cost *C* (Equation 2; represented in the Actor within the dACC_Action_). The dynamical form of these equations is described in the *dACC*_*Boost*_*- LC-VTA system* paragraph in Supplementary Methods.

### LC

#### LC control over effort exertion and behavioural activation

The LC module plays a double role (Figure 1c, orange box). First it controls cost via parameter *Ne*, as a function of boosting value *b* selected by the dACC_Boost_ module. For sake of simplicity, we assigned to *Ne* the value of *b*; any monotonic function would have played a similar role.

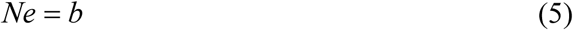

The *Ne* signal is directed also toward external brain areas as a performance modulation signal (Figure 2; Simulation 2b).

#### LC control over learning rate

The LC module optimizes also learning rate in the two dACC modules (*λ* and *λ*_*B*_). Approximate optimization of *λ* solves the trade-off between stability and plasticity, increasing learning speed when the environment changes and lowering it when the environment is simply noisy. In this way, the RML updates its knowledge when needed (plasticity), protecting it from random fluctuations. This function is performed by means of recurrent connections between the dACC (both modules) and the LC module, which controls learning rate based on the signals afferent from the dACC. The resulting algorithm approximates Kalman filtering (Kalman, 1960; Welch and Bishop, 1995), which is a recursive Bayesian estimator. In its simplest formulation, Kalman filter computes expectations (posteriors) from current estimates (priors) plus PE weighted by an adaptive learning rate (called Kalman gain). If we define process variance as the outcome variance due to volatility of the environment, Kalman filter computes the Kalman gain as the ratio between process variance and total variance (i.e. the sum of process and noise variance). From the Bayesian perspective, the Kalman gain reflects the confidence about priors, so that high values reflect low confidence in priors and more influence of evidence on posteriors estimation.

The main limitation of this and similar methods is that one must know a priori the model describing the environment statistical properties (noise and process variance). This information is typically inaccessible by biological or artificial agents, which perceive only the current state and outcome signals from the environment. Our LC module bypasses this problem by an approximation based on the information afferent from the dACC, without knowing a priori neither process nor noise variance. To do that, the LC modulates *λ* (or *λ_B_*) as a function of the ratio between the estimated variance of state/action-value (V âr (v)) over the estimated squared PE 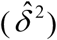:

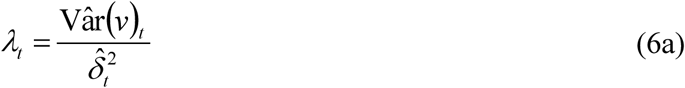

With process variance given by:

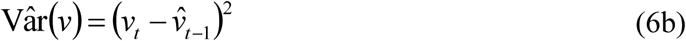

Where 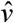 is the estimate of *v*, obtained by low-pass filtering tuned by meta-parameter *α*:

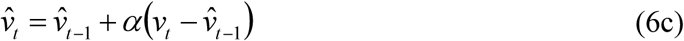

The same low-pass filter is applied to PE signal (*δ*) to obtain a running estimation of total variance 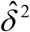, which corresponds to the squared estimate of unsigned PE:

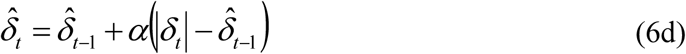

For numerical stability reasons, *λ* was constrained in an interval between 1 (max learning rate) and *β* (min learning rate).

In summary, in equations 6a-d Kalman gain is approximated using 3 components: reward expectation (*v*), PE signals (*δ*) (both afferent from the dACC modules) and a meta-parameter (*α*), defining the low-pass filter to estimate process and total variance. The meta-parameter *α* represents the minimal assumption that noise-related variability occurs at a faster time scale than volatility-related variability. Equations 6a-d are implemented independently for each of two dACC modules, so that each Critic interacts with the LC to modulate its own learning rate. The dACC modules and the LC play complementary roles in controlling *λ*: The dACC modules provide the LC with the time course of expectations and PEs occurring during a task, while the LC integrates them to compute Equation 6a.

The dynamical form of these equations is described in the *dACC-LC system* paragraph in Supplementary Methods.

### VTA

The VTA provides training signal *DA* to both dACC modules, either for action selection directed toward the environment (by dACC_Action_) or for boosting-level selection (by dACC_Boost_) directed to the midbrain catecholamine nuclei (Figure 1c, red box). The VTA module also learns to link dopamine signals to arbitrary environmental stimuli (non-primary rewards) to allow higher-order conditioning. We hypothesize that this mechanism is based on DA shifting from primary reward onset to conditioned stimulus (*s*, *a*, or both) onset (Ljungberg et al., 1992).

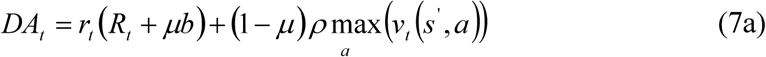

Equation 7a represents the modulated (by *b*) reward signal. Here, *r* is a binary variable indicating the presence of reward signal, and *R* is a real number variable indicating reward magnitude. Parameter *ρ* is the TD discount factor, while parameter *μ* is a scaling factor distributing the modulation *b* between primary (first term of the equation) and non-primary (second term) reward. It is worth noting that when *μ* = 0, Equation 7a simplifies to a Q-learning reward signal.

The VTA signal directed toward the dACC_Boost_ is described by the following equation:

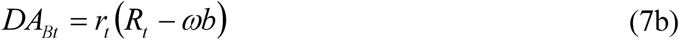

where *ω* is a parameter defining the cost of catecholamine boosting (Kool et al., 2010; Kool and Botvinick, 2013; Shenhav et al., 2013). In summary, boosting up DA by *b* (Equation 7a), can improve behavioural performance (as shown in simulations below) but it also represents a cost (Equation 7b). The dACC_Boost_ module finds the optimal solution for this trade-off, choosing the optimal DA level to maximize performance while minimizing costs (for a formal analysis about this optimization process we refer to Verguts et al., 2015). The dynamical form of these equations is described in the *dACC*_*Action*_*-VTA* and *dACC*_*Boost*_*-VTA* paragraphs in Supplementary Methods.

### Control over other brain areas

Finally, the RML can optimize performance of other brain areas. It does so via the LC-based control signal (*Ne*), which is the same signal that modulates costs (Equation 2; Figure 2). Indeed, the Actor-Critic function of the dACC_Action_ module is domain-independent (i.e. the state/action channels can come from any brain area outside dACC), and this allows a dialogue with other areas. Moreover, because optimization of any brain area improves behavioural performance, the dACC_Boost_ can modulate (via LC signals) any cortical area to improve performance (see Simulation 2c)

## Results

In this section, we report the results obtained with the dynamical model. Simulation results from the discrete model (reported in paragraph “Supplementary Results: main results replication with discrete model) are equivalent and replicated the results described here. Simulation 2c (related to WM task) required intra-trial dynamics, and was therefore run only with the dynamical version.

To mimic standard experimental paradigms as closely as possible, we repeated just 12 times each simulation (simulated subjects), to test whether the model could generate a large effect size of results. Obviously, p-values improved (but not the effect sizes) when running more simulated subjects.

### Simulation 1: learning rate and Bayesian inference

Adaptive control of learning rate is a fundamental aspect of cognition. Humans can solve the tradeoff between stability and plasticity in a Bayesian fashion, by changing the learning rate as a function of environmental changes (Behrens et al., 2007; Yu, 2007), and distinguishing between variability due to noise versus variability due to actual changes of the environment (Yu and Dayan, 2005; Silvetti et al., 2013a).

We will investigate not only whether the model can capture and explain human adaptive control of learning rate at behavioural level, but also a set of experimental findings at neural level, which have not yet been reconciled in one single theoretical framework. According to these findings, LC activity (and thus Ne release) tracks volatility (probably controlling learning rate) while dACC activation tracks global environmental uncertainty (Nassar et al., 2012; Silvetti et al., 2013a, 2013b).

#### Simulation methods

We administered to the RML a 2-armed bandit task in three different stochastic environments (Figure 3a-b). The three environments were: stationary environment (Stat, where the links between reward probabilities and options were stable over time, either 70 or 30%), stationary with high uncertainty (Stat2, also stable reward probabilities, but all the options led to a reward in 60% of times), and volatile (Vol, where the links between reward probabilities and options randomly changed over time) (see also Table s2). We assigned higher reward magnitudes to choices with lower reward probability, to promote switching between choices and to make the task more challenging (cf. Behrens et al. 2007). Nonetheless, the value of each choice (probability × magnitude) remained higher for higher reward probability (see Supplementary Methods for details), meaning that reward probability was the relevant variable to be tracked. A second experiment, where we manipulated reward magnitude instead of reward probability (see: Supplementary Results: dynamical model; Figure s7), led to very similar results.

**Figure 3.**
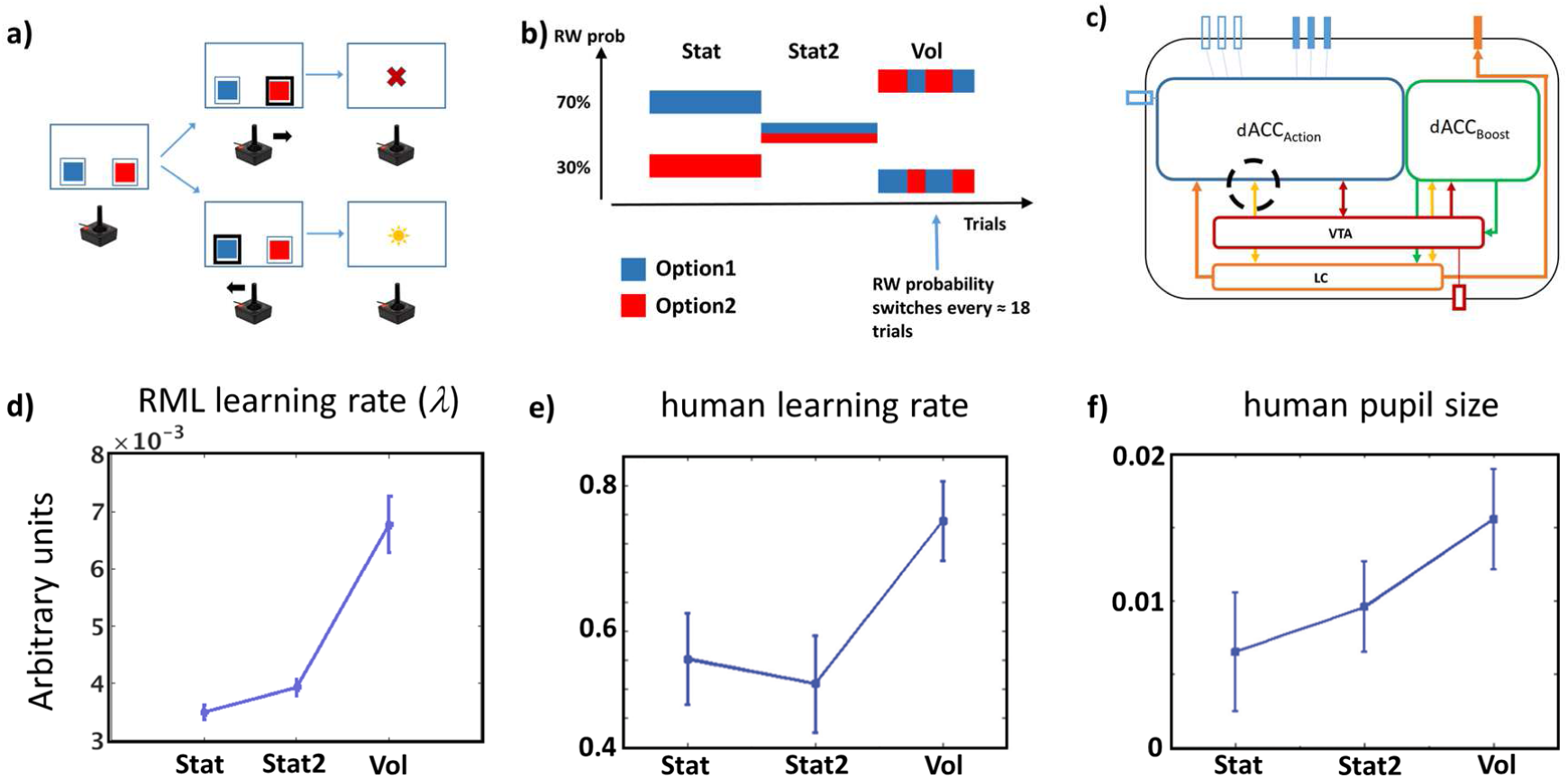
**a)** The task (2-armed bandit) is represented like a binary choice task (blue or red squares), where the model decisions are represented as joystick movements. After each choice, the model received either a reward (sun) or not (cross). **b)** Example of time line of statistical environments (order of presentation of different environments was randomized across simulations). The plot shows reward probability linked to each option (blue or red) as a function of trial number. In this case the model executed the task first in a stationary environment (Stat), then in a stationary environment with high uncertainty (Stat2), and finally in a volatile (Vol) environment. **c)** Model schema showing where we recorded the signal to measure the learning rate variation (dashed black circle). **d)** Learning rate as a function of environmental volatility (± s.e.m.) in the RML and humans **e)** (modified from: Silvetti et al., 2013a). **f)** human LC activity (inferred by pupil size; Joshi et al. 2016; Varazzani et al. 2015; Aston-Jones and Cohen 2005) during the same task.

#### Simulation Results and Discussion

The RML performance in terms of optimal choice percentages was: Stat = 66.5% (± 4% s.e.m.), Vol = 63.6% (± 1.4% s.e.m.). For Stat2 condition there was no optimal choice, as both options led to reward in 60% of times. Importantly, the model successfully distinguished not only between Stat and Vol environments, but also between Stat2 and Vol, increasing the learning rate exclusively in the latter (Figure 3d). There was a main effect of volatility on learning rate *λ* (F(2,11) = 29, p < 0.0001). Post-hoc analysis showed that stationary conditions did not differ (Stat2 > Stat, t(11) = 1.65, p = 0.13), while in volatile condition learning rate was higher than in stationary conditions (Vol > Stat2, t(11) = 5.54, p < 0.0001; Vol > Stat, t(11) = 5.76, p < 0.0001). Hence, interaction between dACC and LC allows disentangling uncertainty due to noise from uncertainty due to actual changes (Yu and Dayan, 2005; Silvetti et al., 2013a), promoting flexibility (high learning rate) when new information must be acquired, and stability (low learning rate) when acquired information must be protected from noise. This mechanism controls learning rates in both the dACC_Action_ and the dACC_Boost_ modules, thus influencing the whole RML dynamics. The same learning rate effect was found in experimental data (Figure 3e). Indeed, humans increased both learning rate and LC activity only in Vol environments (Silvetti et al., 2013). Thus, humans could distinguish between outcome variance due to noise (Stat2 environment) and outcome variance due to actual environmental change (Vol environment).

The model was also consistent with the fMRI data cited above (Silvetti et al., see Figure 4). During a RL task executed in the same three statistical environments used in this simulation, the human dACC activity did not follow the pattern in Figure 3d but instead peaked for Stat2 environment, suggesting that activity of human dACC is dominated by prediction error operations rather than by explicit estimation of environmental volatility (Figure 4a). The RML captures these results (Figure 4b) by its PE activity. Finally, it is worth noting that Vol-related activity of both model and human dACC was higher than in Stat environment (stationary with low uncertainty), thus replicating the results of previous fMRI studies by Behrens et al. (2007).

**Figure 4.**
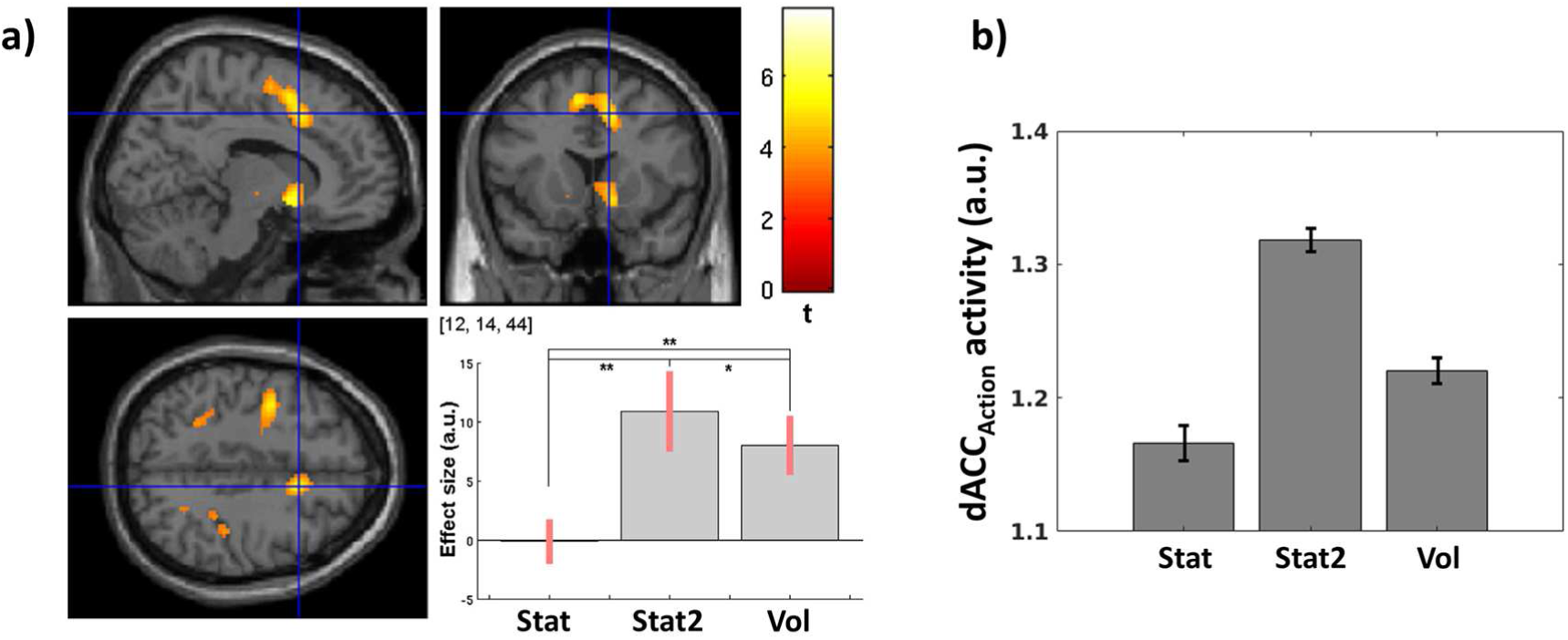
**a)** dACC activity effect size (extracted from the ROI indicated by cross) in a RL task executed during fMRI scanning. The task was performed in the same three environments we used in our simulations. dACC activity peaked in Stat2 and not in Vol condition (modified from: Silvetti et al., 2013b). **b)** dACC_Action_ average prediction error activity (sum of *δ* units activity ± s.e.m.) as a function of environmental uncertainty. Differently from the LC, the dACC is maximally active in stationary uncertain environments (Stat2).

Summarizing, results in Figure 3d show the effectiveness of our RL-based Kalman approximation, suggesting that the LC codes for volatility and that our algorithm modeling dACC-LC dialogue is both computationally effective and neurophysiologically grounded. Results in Figure 4b, on the other hand, indicate that, although the dACC is part of a Bayesian estimator, its activity is mostly influenced by overall environmental variance, rather than coding for volatility.

### Simulation 2: Controlling physical and cognitive effort

A long list of experimental results indicates DA and Ne neuromodulators as crucial not only for learning environmental regularities, but also for exerting cognitive control (e.g. Aston-Jones & Cohen 2005; Sara 2009; Vijayraghavan et al. 2007; Langner & Eickhoff 2013; D’Esposito & Postle 2015). Although these mechanisms have been widely studied, only few computational theories explain how the midbrain catecholamine output is controlled to maximize performance (Doya, 2002; Yu and Dayan, 2005; Niv et al., 2007), and how the dACC is involved in such a process. In this section, we describe how the dACC_Boost_ module learns to regulate LC and VTA activity to control effort exertion, at both cognitive and physical level (Chong et al., 2017), to maximize long-term reward. In Simulation 2a, we test the cortical-subcortical dynamics regulating catecholamine release in experimental paradigms involving decision-making in physically effortful tasks, where cost/benefit trade off must be optimized (Salamone et al., 1994). In Simulation 2b, we show how the LC can provide a Ne signal to external “client” systems to optimize cognitive effort allocation and thus behavioural performance in a visuo-spatial WM task. In both simulations, we also test the RML dynamics and behaviour after DA lesion.

#### Simulation 2a: Physical effort control and decision-making in challenging cost/benefit trade off conditions

Deciding how much effort to invest to obtain a reward is crucial for human and non-human animals. Animals can choose high effort-high reward options when reward is sufficiently high. The impairment of the DA system strongly disrupts such decision-making (Salamone et al., 1994; Walton et al., 2009). Besides the VTA, experimental data indicate also the dACC as having a pivotal role in decision-making in this domain (Kennerley et al., 2011; Apps and Ramnani, 2014; Vassena et al., 2014). In this simulation, we show how cortical-subcortical interactions between the dACC, VTA and LC can drive optimal decision-making when effortful choices leading to large rewards compete with low effort choices leading to smaller rewards. We thus test whether the RML can account for both behavioral and physiological experimental data. Moreover, we test whether simulated DA depletion in the model can replicate the disruption of optimal decision-making, and, finally, how effective behaviour can be restored.

##### Simulation Methods

We administered to the RML a 2-armed bandit task with one option requiring high effort to obtain a large reward, and one option requiring low effort to obtain a small reward (Walton et al. 2009; here called Effort task; Figure 5a). The task was also administered to a DA lesioned RML (simulated by reducing all the outputs from VTA module, see Experimental Methods section in Supplementary Matherial).

**Figure 5.**
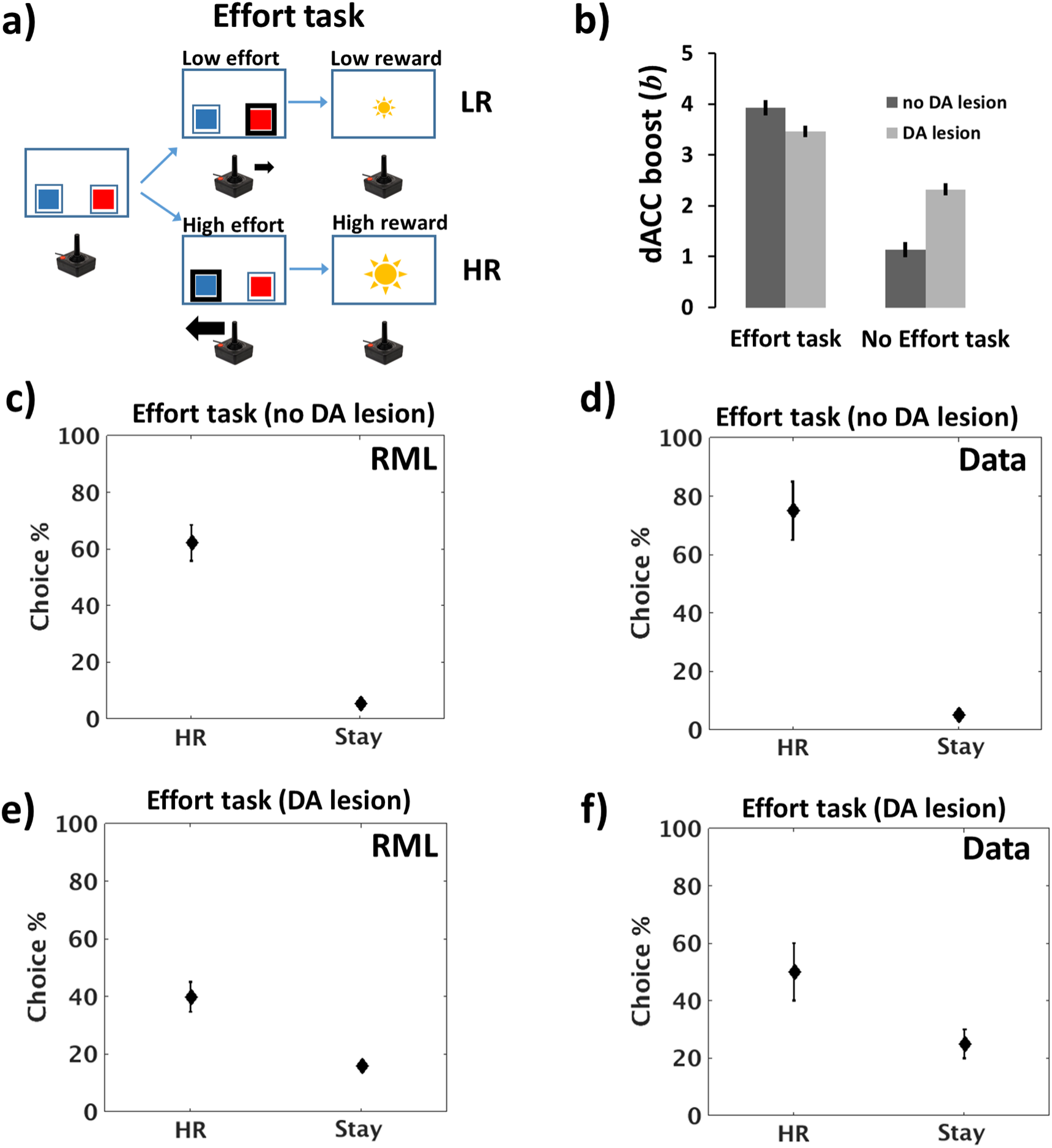
**a)** Effort task, where a high effort choice (thick arrow from joystick) resulting in high reward (HR, large sun) was in competition with a low effort choice (thin arrow) leading to low reward (LR, small sun). **b)** Catecholamines boosting (*b*) as a function of task type (Effort or No Effort task) and DA lesion. The boosting value (recorded from the decision units within the dACC_Boost_ module) is higher in the Effort task (main effect of task), but there is also a task x lesion interaction indicating the dACC_Boost_ attempts to compensate the loss of DA, to achieve at least LR (see main text). **c)** Behavioural results (average HR/(LR+HR) ratio ±s.e.m., and average Stay-to-total choices ratio percentage ±s.e.m.) from RML and **d)** empirical data. **e)** Behavioural results after DA lesion in RML and **f)** in empirical data. In this case animals and the RML switch their preference toward the LR option (requiring low effort). In both d) and f), animal data are from Walton et al. (2009).

Like in Walton et al. (2009), before the execution of the Effort task, the RML learned the reward values in a task where both options implied low effort (No Effort task). Besides the high effort and low effort choices, the model could choose to execute no action if it evaluated that no action was worth the reward (“Stay” option).

##### Simulation Results and Discussion

As shown in Figure 5b, the dACC_Boost_ increased the boosting level (*b*) in the Effort task (main effect of task, F(1,11) = 231.73, p < 0.0001) enhancing both LC and VTA output (equations 5, 7a). Increased *Ne* influences the Actor of the dACC_Action_ (effect of *Ne* on action cost estimation in decision-making process, Equation 2), facilitating effortful actions, while increased DA affects the learning process of the dACC_Action_ (equations 1, 7a), increasing the reward signal related to effortful actions (therefore “subjectively” increasing their value with respect to the “objective” reward signal provided by the environment). At the same time, the dACC_Boost_ had to face the cost of boosting (*ωb* term in Equation 7b), so that the higher *b*, the higher was the reward discount for the dACC_Boost_ module. The result of these two opposite forces (maximizing performance by catecholamines boosting and minimizing the cost of boosting itself) converges to the optimal value of *b* and therefore of catecholamines release by VTA and LC (Figure 6a).

**Figure 6.**
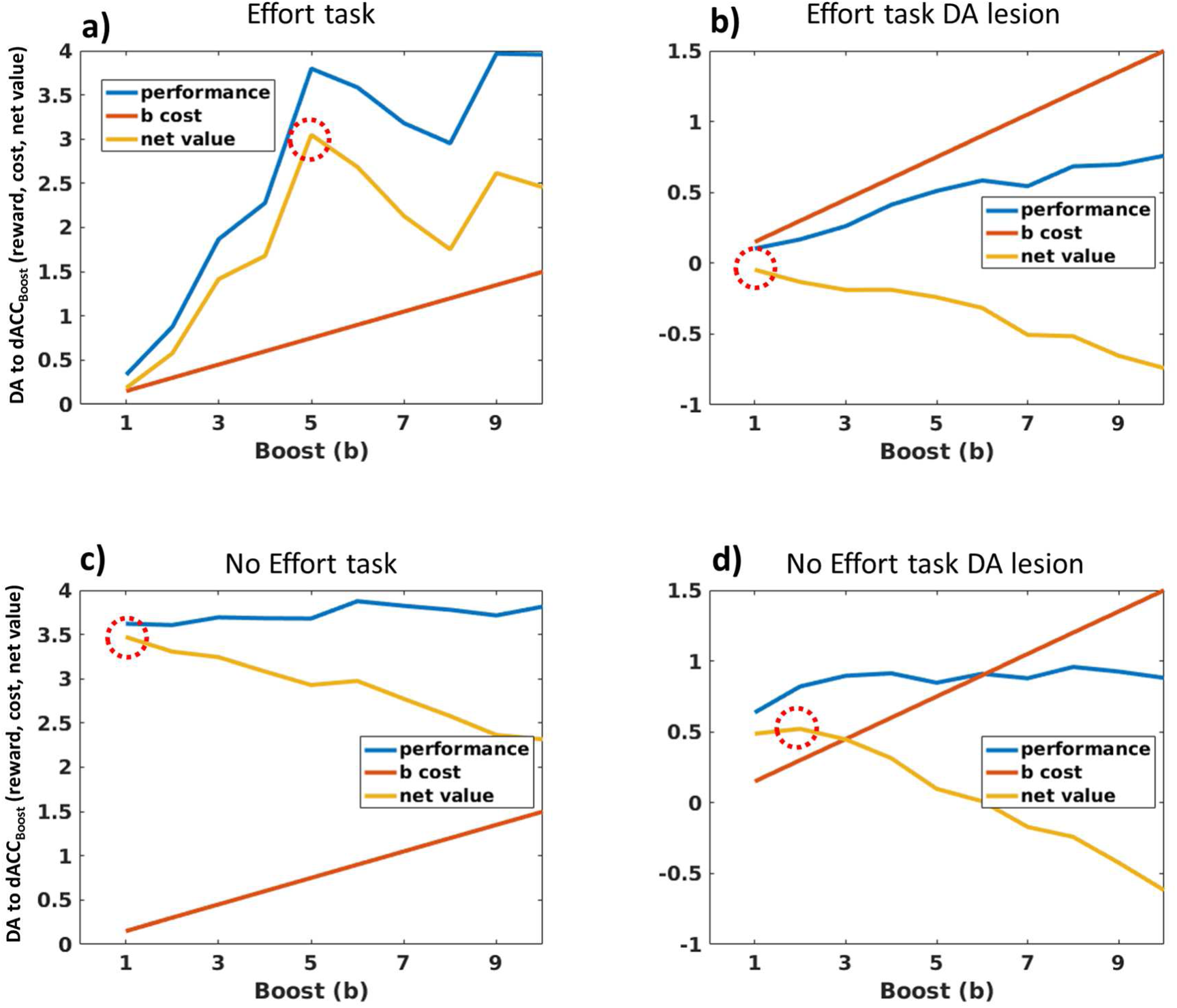
Cost-benefits plots and optimal control of *b* in the dACC_Boost_ module. To obtain these plots we systematically clamped *b* at several values (from 1 to 10, x axis of each plot) and then we administered the same paradigms of Figure 5b (all the combinations Effort x DA lesion). In all the pots, y axis represents simultaneously behavioural performance in terms of average reward (for blue plots), boosting cost (red plots) and net value (performance – boost cost, in other words, Equation 7b). To minimize computation time, we use for these simulations only the discrete model. a) **Effort task, no lesion**. Plot showing RML behavioural performance as a function of *b* (blue plot), boosting cost (red plot, *ωb* member in Equation 7b) and net value for the dACC_Boost_ module (yellow plot, resulting from Equation 7b). Red dotted circles highlight the optimal *b* value which maximizes the final net reward signal received by the dACC_Boost_ module. **b) Effort task, DA lesion.** Same as a, but in this case the RML was DA lesioned. Due to lower average reward signal (blue plot), the net value plot (yellow) is much lower than in a), as the cost of boosting (red plot) did not change. Red dotted circle highlights the optimal *b* value, which is lower than in a). It must be considered that, despite the optimal *b* value is 1, the average *b* (as shown in figures 5b and s11b) is biased toward higher values, as it is selected by a stochastic process (Equation 4) and values lower than 1 are not possible (asymmetric distribution). **c) No effort task, no lesion.** In this case, being the task easy, the RML reaches a maximal performance without high values of *b*, therefore the optimal *b* value is low also in this case. **d) No effort task, DA lesion.** As shown also in Figure 5b, in this case the optimal *b* value (dotted circle), is higher than in c), because a certain amount of boosting is necessary to avoid apathy (moving as an intrinsic cost) and it can give access to large rewards. Apathy is determined by the RML preference for “Stay” option, which gives no reward but it has no costs, boosting can help avoiding apathy.

After DA lesion, the dACC_Boost_ decreased the boosting output during the Effort task, while it increased the boosting output during the No Effort task (task x lesion interaction F(1,11) = 249.26, p < 0.0001). Decreased boosting derives from decreased DA signal to dACC_Boost_ module (Figure 6b). On the other side, increased boosting *b* in No Effort task can be interpreted as a compensatory mechanism ensuring the minimal catecholamines level to achieve the large reward when just a low effort is necessary (Figure 6d). Indeed, the lack of compensation in No Effort task would result in a policy mimicking behavioral apathy, where the RML would often select the “Stay” action, saving on minimal costs of moving but also reducing the amount of reward. in other words, when the incentive is high (high reward available) and the effort required to obtain the reward is low, the RML predicts that the DA lesioned animal would choose to exert some effort (boosting up the remaining catecholamines) to promote active behaviour versus apathy.

At behavioural level, in the Effort task, the RML preferred the high effort option to get a large reward (Figure 5c; t(11) = 4.71, p = 0.0042). After the DA lesion, the preference toward high effort-high reward choice reversed (Figure 5d; t(11) = - 3.71, p = 0.0034). Both results closely reproduce animal data (Walton et al., 2009). Furthermore, the percentage of “Stay” choices increased dramatically (compare figures 5c and 5e; t(11) = 18.2, p < 0.0001). Interestingly, the latter result is also in agreement with animal data and could be interpreted as a simulation of apathy due to low catecholamines level.

#### Simulation 2b: performance recovery after DA lesion, in cost/benefit trade off conditions

In DA lesioned subjects, the preference for HR option can be restored by removing the difference in effort between the two options (Walton et al., 2009), in other words by removing the critical trade-off between costs and benefits. In these simulations, we show how the RML can recover a preference toward HR options when exposed to the same experimental paradigms used in rats.

##### Simulation Methods

The same DA lesioned subjects of Simulation 2a were exposed to either a No Effort task (where both the option required low effort) or a Double Effort task (where both the options required a high effort) (Figure 7a-b). All other experimental settings were identical to those of Simulation 2a.

**Figure 7.**
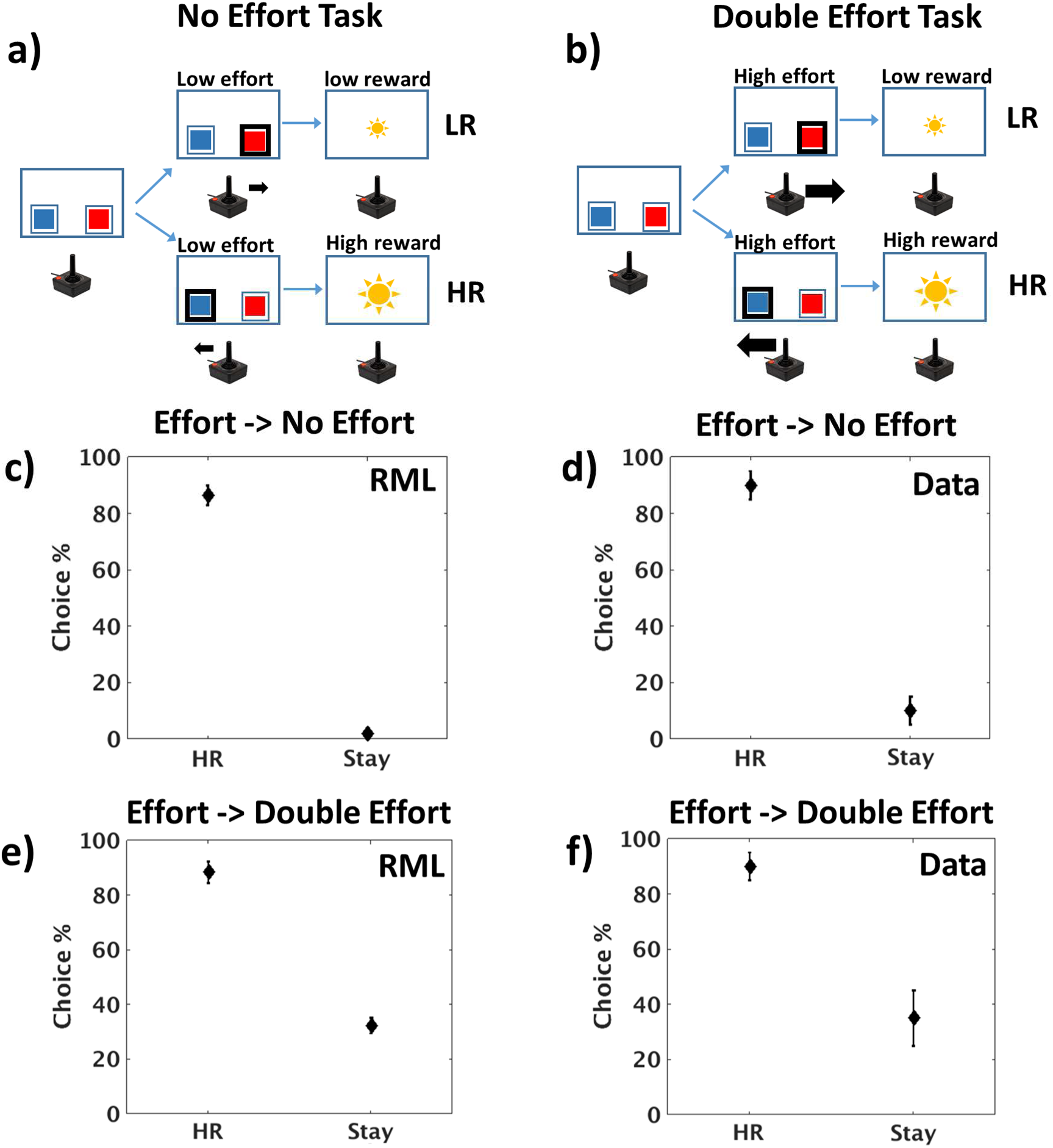
Recovery of HR option preference in DA lesioned subjects. **a)** No Effort task, consists in two possible choices, both requiring low effort to be executed (small black arrow), one leading to a high reward (large sun; HR), the other to a low reward (LR). **b)** Double Effort task, where both options implied high effort. **c-d)** Recovery of the preference for HR option (HR/(HR+LR)) when a No Effort task is administered after an Effort task session (Effort → No Effort), in both RML and animals (mean percentage ± s.e.m.). **e-f)** Same phenomenon when a Double Effort session follows an Effort one (Effort → Double Effort), in both RML and animals. Note that in this case the amount of “Stay” choices (Stay/number of trials) increased, simulating the emergence of apathic behaviour. Animal data from Walton et al., 2009.

##### Simulation Results and Discussion

DA-lesioned RML performance recovers immediately when a No Effort task is administered after the Effort task (Figure 7c), in agreement with animal data by Walton et al. (2009; Figure 7d). This result shows that performance impairment after DA lesion is not due to learning deficit (although partial learning impairment must occur due DA role in learning), but rather to down-regulation of catecholamines boosting, driven by dACC_Boost_. A task where both the options require a low effort does not require a strong behavioural energization, therefore the information about the high reward location is sufficient for an optimal execution.

The same performance recovery occurs also in a task where both options are effortful (Double Effort task, Figure 7e), again in agreement with experimental data (Figure 7f). Also in this case, when there is no trade-off between costs and benefits (both the options are the same in terms of effort), the information about the high reward location is sufficient to correctly execute the task, although there is a reduced catecholamine boosting. Nonetheless, differently from the previous scenario, here emerges a new phenomenon: *apathy* (percentage of “Stay”, Figure 7e-f). Indeed, both the RML and animals often refuse to engage in the task, and rather than working hard to get the high reward (whose position is well known) they prefer to remain still. Apathic behaviour in this experiment is more evident than in Figure 5e-f, because both RML and animals are forced to make an effort to get a reward, while in Simulation 2a (Figure 5e-f) they could opt for the low effort-low reward choice.

#### Simulation 2c: Adapting cognitive effort in a WM task

Ne neuromodulation plays also a crucial role in WM, improving signal to noise ratio by gain modulation mediated by *α*2-A adrenoceptors (Aston-Jones and Cohen, 2005; Wang et al., 2007), and low level of Ne transmission leads to WM impairment (Li and Mei, 1994; Li et al., 1999). At the same time, as described above, it is a major biological marker of effort exertion (Kahneman, 1973; Varazzani et al., 2015). Besides Ne release by the LC, experimental findings showed that also dACC activity increases as a function of effort in WM tasks, e.g. in mental arithmetic (Borst and Anderson, 2013; Vassena et al., 2014). Here we show that the same machinery that allows optimal physical effort exertion (Simulation 2a) may be responsible for optimal catecholamine management to control the activity of other brain areas, thus rooting physical and cognitive effort exertion in a common decision-making mechanism. This is possible because the design of the RML allows easy interfacing with external systems. Stated otherwise, the macro-circuit dACC-midbrain works as a “server” providing control signals to “client” areas to optimize their function. Given the dynamical nature of this simulation (we used a dynamical model of WM, and the task implies an intra-trial delay when information must be retained), we used in this case only the dynamical version of the RML.

##### Simulation Methods

We connected the RML to a WM model (FROST model; Ashby et al. 2005; see “FROST model description” section in Supplementary Methods). Information was exchanged between the two models through the state/action channels in the dACC_Action_ module and the external LC output. The FROST model was chosen for convenience only; no theoretical assumptions prompted us to use this model specifically. FROST is a dynamical recurrent neural network simulating a macro-circuit involving the DLPFC, the parietal cortex and the basal ganglia. This model simulates behavioural and neurophysiological data in several visuo-spatial WM tasks. FROST dynamics simulates the effect of memory loads on information coding, with a decrement of coding precision proportional to memory load (i.e. the number of spatial locations to be maintained in memory). This feature allows to simulate the increment of behavioural errors when memory load increases (Ashby et al., 2005). The external LC output (*Ne*) improves the signal gain in the FROST DLPFC neurons, increasing the coding precision of spatial locations retained in memory (Equation s21 in Supplementary Methods), thus improving behavioural performance. We administered to the RML-FROST circuit a delayed matching-to-sample task with different memory loads (a template of 1, 4 or 6 items to be retained; Figure 8a). We used a block design, where we administered three blocks of 70 trials, each with one specific memory load (1, 4, or 6). In 50% of all trials, the probe fell within the template. The statistical analysis was conducted by a repeated measure 3x2 ANOVA (memory load and DA lesion).

**Figure 8.**
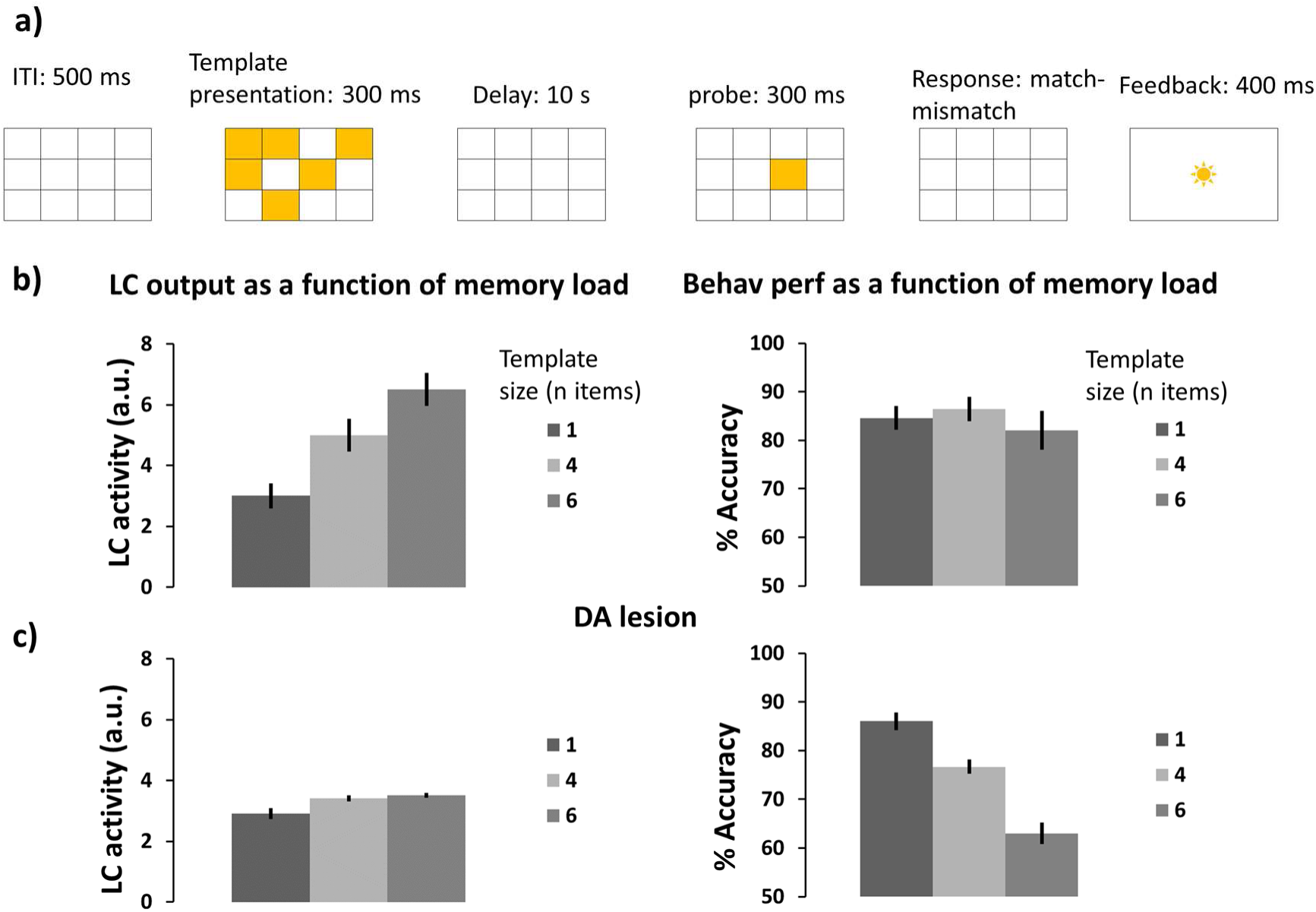
**a)** Delayed Matching-to-sample task: events occurring in one trial. **b)** Left: LC activity (*Ne* signal) as a function of memory load (number of items presented in the template). Right: behavioural performance as a function of memory load. **c)** LC activity (*Ne* signal) and behavioural performance after DA lesion. Error bars indicate ±s.e.m.

##### Simulation Results and Discussion

The dACC_Boost_ module dynamically modulates *Ne* release as a function of memory load, in order to optimize performance (Figure 8b, left panel; main effect of memory load on LC output: F(2,22)= 16.74, p < 0.0001). Like in Simulation 2a, in case of DA lesion, the VTA-dACC-LC interaction is disrupted, leading to a devaluation of boosting and the consequent decision (by the dACC_Boost_ module) of downregulating LC activity (Figure 8c, left panel; main effect of DA lesion on LC output: F(1,11)= 24.88, p < 0.0001). This happened especially for high memory loads (lesion × memory-load interaction: F(2,22) = 7.1, p = 0.0042). LC modulation impairment results in poor performance in particular for high memory loads, when high level of *Ne* is necessary (Figure 8c, accuracy, right panel; lesion × memory-load interaction: F(2,22) = 8.6, p = 0.0017).

### Simulation 3: Reinforcement Learning, meta-learning and higher-order conditioning

Animal behavior in the real world is seldom motivated by conditioned stimuli directly leading to primary rewards. Usually, animals have to navigate through a problem space, selecting actions to come progressively closer to a primary reward. In order to do so, animals likely exploit both model-free and model-based learning (Niv et al., 2006; Pezzulo et al., 2013; Walsh and Anderson, 2014). Nonetheless, model-free learning from non-primary rewards (i.e. higher-order conditioning) remains a basic key feature for fitness, and the simplest computational solution to obtain adaptive behaviour in complex environments. For this reason, we focus on model-free learning here.

A unifying account explaining behavioral results and underlying neurophysiological dynamics of higher-order conditioning is currently lacking. First, at behavioral level, literature suggests a sharp distinction between higher-order conditioning in classical versus instrumental paradigms. Indeed, although it is possible to train animals to execute complex chains of actions to obtain a reward (instrumental higher-order conditioning, Pierce and Cheney, 2004), it is impossible to install a third- or higher-order level of classical conditioning (i.e. when no action is required to get a reward; Denny and Ratner, 1970). Although the discrepancy has been well known for decades, its reason has not been resolved. Second, a number of models have considered how TD signals can support conditioning and learning more generally (Holroyd and Coles, 2002; Williams and Dayan, 2005). However, no model addressing DA temporal dynamics also simulated higher-order conditioning at behavioural level.

Here we use the RML to provide a unified theory to account for learning in classical and instrumental conditioning. We show how the RML can closely simulate the DA shifting in classical conditioning (Figure s8 in: “Supplementary Results: dynamical model”). We also describe how the VTA-dACC interaction allows the model to emancipate itself from primary rewards (higher-order conditioning). Finally, we investigate how the synergy between the VTA-dACC_Boost_ and LC-dACC_Boost_ interactions (the catecholamines boosting dynamics) is necessary for obtaining higher-order instrumental conditioning. This provides a mechanistic theory on why higher-order conditioning is possible only in instrumental and not in classical conditioning.

### Simulation 3a: Higher-order classical conditioning

Equation 7a (and its dynamical homologous Equation s5b) expresses a progressive linking of DA response to conditioned stimuli. This is due to the max(v()) term in Equation 7a and the time derivative of *v* in Equation s5b for the dynamical version of RML. As VTA can vigorously respond to conditioned stimuli, it is natural to wonder whether a conditioned stimulus can work as a reward itself, allowing to build a chain of progressively higher-order conditioning (i.e. not directly dependent on primary reward). However, for unknown reasons, classical higher-order conditioning is probably impossible to obtain in animal paradigms (Denny and Ratner, 1970; O’Reilly et al., 2007). We thus investigate what happens in the model in such a paradigm.

#### Simulation Methods

We first administered a first-order classical conditioning. We then conditioned a second cue by using the first CS as a non-primary reward. The same procedure was repeated up to third-order conditioning. Each cue was presented for 2s followed by the successive cue or by a primary reward. All cue transitions were deterministic and the reward rate after the third cue was 100%.

#### Simulation Results and Discussion

In Figure 9 we show the VTA response locked to the onset of each conditioned stimulus. Surprisingly, but in agreement with experimental animal data, the conditioned cue-locked DA release is strongly blunted at the 2^nd^ order, and disappeared almost completely at the 3^rd^ order. This aspect of VTA dynamics is naturally captured by the dynamical version of the RML (closer to neurophysiology than the discrete version), because at each order of conditioning, the cue-locked signal is computed as the temporal derivative of reward prediction unit activity (Equation s5b), losing part of its power at each conditioning step. In the discrete version of the model, this aspect is expressed in Equation 7a, where (like in Q-learning) the cue-locked DA response is scaled by a positive real number smaller than one (*ρ*). This mechanism implies a steep decay of the conditioning effectiveness of non-primary rewards, as at each order of conditioning, the reinforcing property of cues is lower and lower. From the ecological viewpoint, it makes sense that the weaker is the link between a cue and a primary reward, the weaker should be its conditioning effectiveness. Nonetheless, as we describe in the following paragraph, this phenomenon is in some way counteracted in instrumental conditioning.

**Figure 9.**
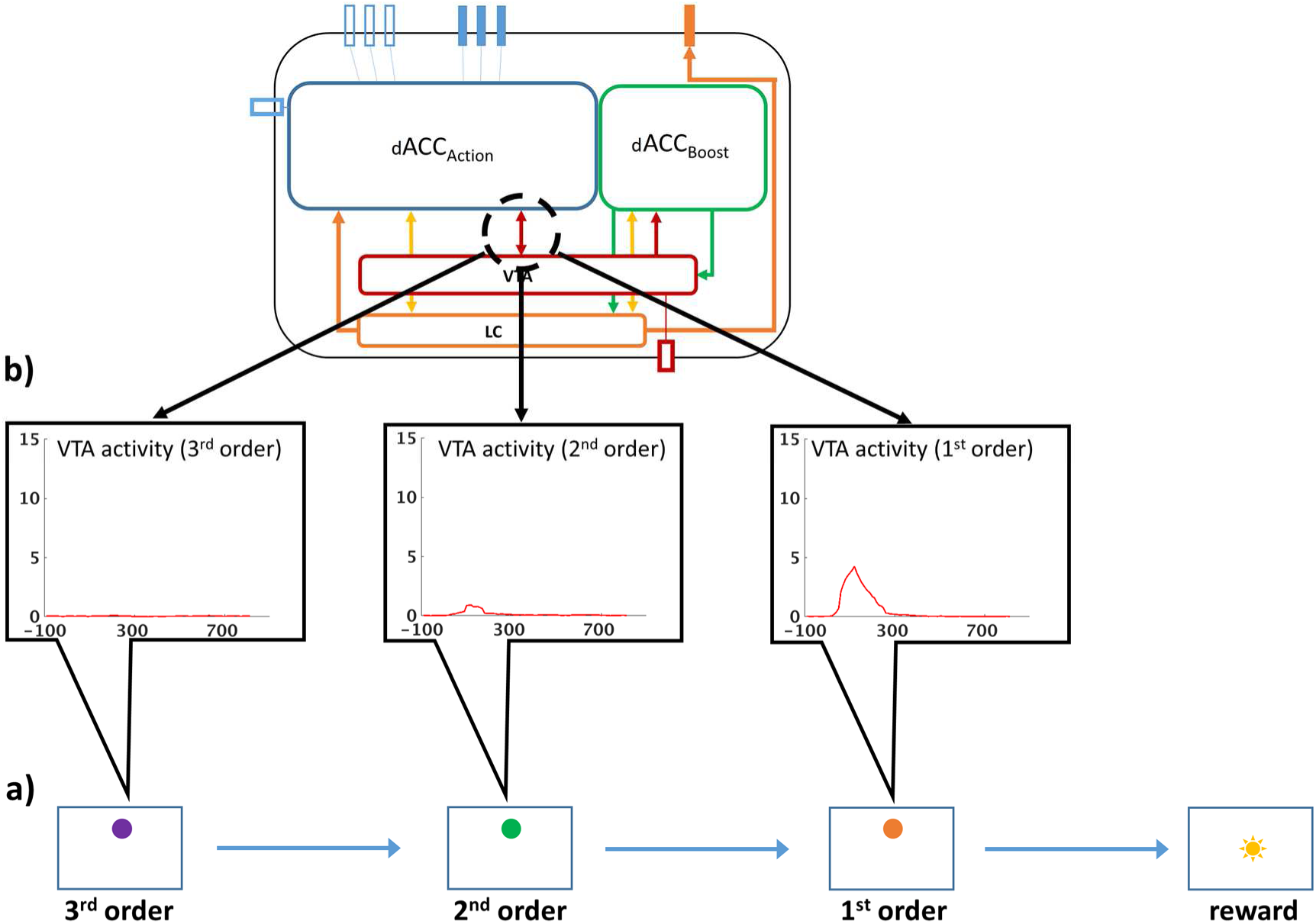
**a)** Experimental paradigm for higher-order classical conditioning. Sequence of conditioned stimuli (colored disks) followed by primary reward (sun). **b)** VTA activity (DA signal) locked to each of conditioned stimuli. Dashed black circle indicates where the plotted signals were recorded from: DA signal from Equation s5b (for RML dynamical version) or from Equation 7a (for RML discrete version).

#### Simulation 3b: Chaining multiple actions and higher-order conditioning

Differently from classical conditioning paradigms, animal learning studies report that in instrumental conditioning it is possible to train complex action chains using conditioned stimuli (environmental cues) as reward proxies, delivering primary reward only at the end of the task (Pierce and Cheney, 2004).

##### Simulation Methods

We administered to the RML a maze-like problem, structured as a series of binary choices before the achievement of a final reward (Figure s6). Each choice led to an environmental change (encoded by a colored disk, like in Figure 2). The training procedure was the same as for higher-order classical conditioning. We first administered a first-order instrumental conditioning (2-armed bandit task). Then, we used the conditioned environmental cue as non-primary reward to train the RML for second-order conditioning. The procedure was repeated up to third-order conditioning. State-to-state transitions were deterministic and primary reward rate was 100% for correct choices and 0% for wrong choices.

##### Simulation Results and Discussion

At the end of training, the system was able to perform three sequential choices before getting a final reward, for an average accuracy of 77.3% (90% C.I. = ±13%) for the first choice (furthest away from primary reward; purple disk, Figure 10a); 95.8% (90% C.I. = [4.2, 5.6]%) for the second; and 98% (90% C.I. = ±0.4%) for the third choice (the one potentially leading to primary reward; orange disk, Figure 10a). Figure 10b shows the cue-locked VTA activity during a correct sequence of choices. Differently from classical conditioning, the DA signal amplitude persists over several orders of conditioning, making colored disks (also far away from final reward) effective non-primary rewards, able to shape behaviour.

**Figure 10.**
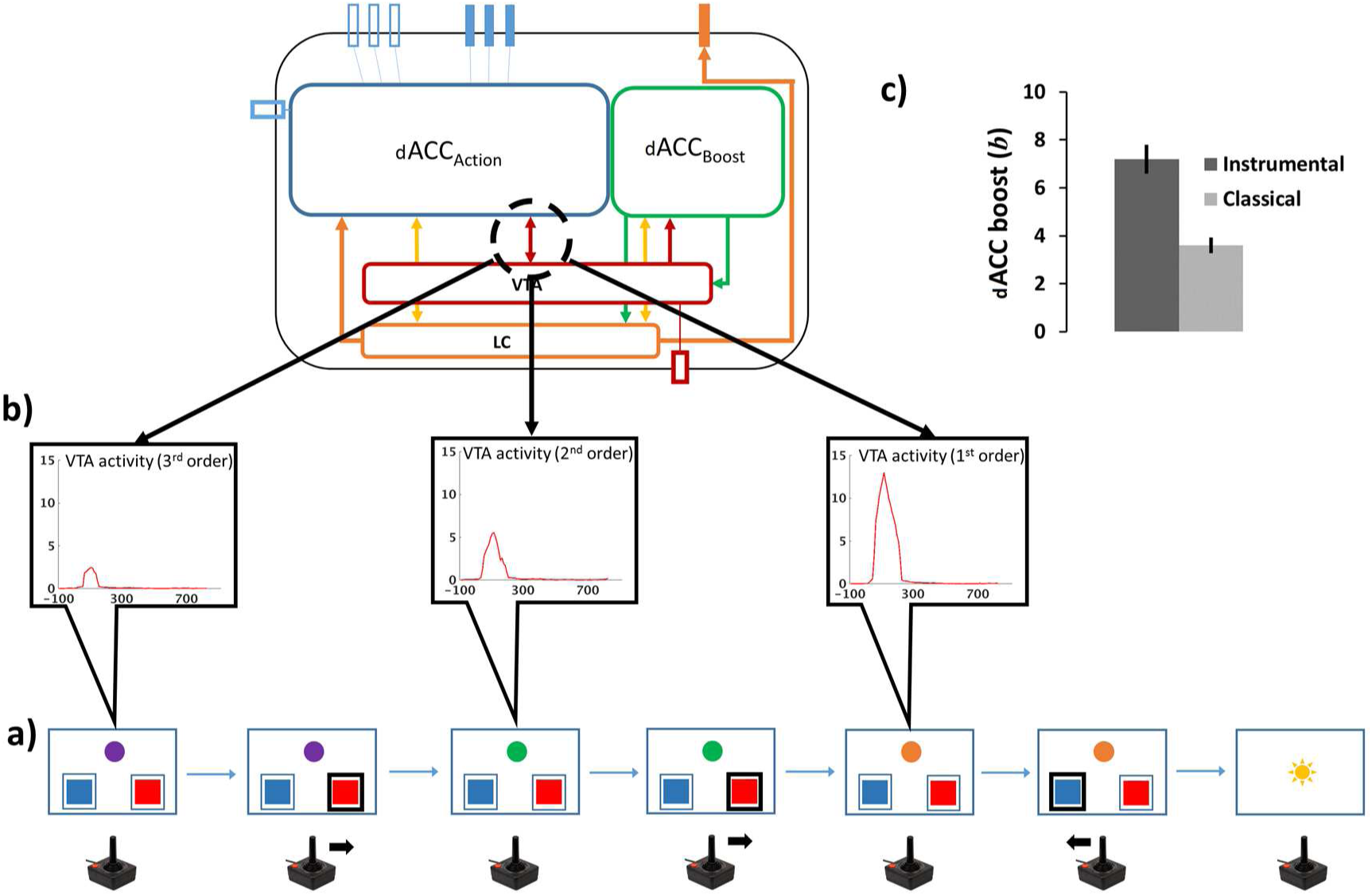
VTA dynamics during higher order instrumental conditioning. **a)** Events occurring during a sequence of correct choices in the task represented also in Figure 2. See Supplementary Methods for details. **b)** Cue-locked (colored disk indicating the environment state) VTA activity. Dashed black circle on the model schema indicates where the plotted signals were recorded from. Differently from higher order classical conditioning, the DA release persists over progressive abstraction of rewards (associative distance from primary reward). **c)** Boosting level (*b*) is higher in instrumental conditioning as compared to classical conditioning (cfr. Figure 9).

The reason for this difference between classical and instrumental conditioning, is in the role played by the dACC_Boost_ module. Figure 10c compares average boosting levels *b* (selected by the dACC_Boost_) in classical and instrumental conditioning. The dACC_Boost_ learned that boosting catecholamines was useful in instrumental conditioning; furthermore it learned that it was not useful in classical conditioning (t(11) = 5.64, p < 0.0001). This decision amplified DA release during task execution only in instrumental conditioning (compare Figure 10b and Figure 9b). Enhanced VTA activity during the presentation of conditioned stimuli (the colored lights indicating a change in the problem space) means more effective higher-order conditioning, therefore a more efficient behaviour. Conversely, in classical conditioning, the model does not need to make any motor decision, as the task consists exclusively of passive observation of incoming cues (colored lights). Therefore, boosting Ne and/or DA does not affect performance (reward amount), as this is completely decided by the environment. In this case, boosting would only be a cost (Equation 7b and its dynamic homologous Equation s18), and the dACC_Boost_ module learned not to boost, with a low DA levels for conditioned stimuli. This explains the strong limitations in establishing higher-order classical conditioning, and shows how decisions about effort exertion are involved in higher-order conditioning.

### General Discussion

We proposed a novel account of the role of dACC in cognition, suggesting that its elusive computational role can be understood by broadening the theoretical view to a recurrent macro-circuit including the catecholaminergic midbrain nuclei. At a theoretical level, this reconciled several previous frameworks of dACC function, including behavioural adaptation from a Bayesian perspective (Kolling et al., 2016), effort modulation for cognitive control (Shenhav et al., 2016), and RL-based action-outcome comparison (Silvetti et al., 2014). Furthermore, the model explained a wide array of heterogeneous empirical data, including learning rate optimization, effort exertion in physical and cognitive tasks, and higher-order classical and instrumental conditioning.

The first meta-learning process we analyzed concerned learning rate (Simulation 1). The RML provides an explicit theory and neuro-computational architecture of how autonomous control of learning rate can emerge from dACC-LC interaction. We propose that the dACC provides signals about environment statistical structure to the LC; in turn, LC estimates optimal learning rate by using Bayesian inference. This explains why both structures are necessary for optimal control of flexibility (Behrens et al., 2007; Nassar et al., 2012; Jepma et al., 2016), and why the dACC activity seems to be related to RL computation (Silvetti et al., 2013b) while the LC activity to volatility estimation (Silvetti et al., 2013a). The RML not only proposes a novel implementation of near optimal learning rate control, but it also unifies RL and Bayesian perspectives, and explains how the mammalian brain can estimate environmental volatility by using RL signals generated during animal-environment interaction (Simulation 1).

The second meta-learning process concerned effort exertion and optimal allocation of cognitive and physical resources to achieve a goal (simulations 2a-c). We proposed that investing effort (cognitive or physical) and controlling associated costs can be seen as an RL problem (Verguts et al., 2015). Differently from earlier models, the RML generalizes this mechanism to any cognitive domain, showing how the dACC-midbrain system could work as a server providing optimal control signals to other brain areas to maximize success while minimizing costs. Moreover, the RML provides an explicit theory about the role of catecholamines in cognitive control and effort exertion. Finally, the modulation of catecholamines release (of which the dACC_Boost_ is a key element) itself is based on Bayesian learning. This aspect provides Bayesian flexibility to cognitive control, a novelty that unifies both these theoretical perspectives.

The third meta-learning process that we simulated concerned control over *reward* signal (both primary and non-primary), introducing for the first time the dimension of subjectivity in RL. The reward signal is no longer an “objective” feedback from the environment, but it can be proactively modulated 1) to increase the value of actions implying effort (thus energizing behaviour; Equation 7a, simulations 2a-c) or 2) to increase the value of non-primary rewards (simulations 3a-b). The latter mechanism, in particular, allowed to explain why higher-order conditioning is possible in instrumental but not in classical paradigms. Moreover, as VTA activity is modulated by the same signal modulating *Ne* release (*b* from dACC_Boost_), this feature provides a unified theoretical view between optimal effort allocation and control over motivational and learning aspects.

In the RML learning rate, effort estimation and reward-related processes are integrated (Figure 1c). For example, dynamic control of learning rate (*λ*) is based on RL signals from dACC modules (*v* and *δ*). Learning rate modulation influences both decision-making for action selection (*a*) and for boosting control (*b*). Boosting control modulates in parallel both LC and VTA, changing both effort investment (*Ne*) and reward signals (*DA*). Catecholamines modulation changes behavioural performance, influencing action selection and environmental feedback, and then influencing back LC control over learning rate.

The RML framework has two main limitations. First, in the RML DA plays a role only in learning. As with any other neuromodulator, experimental results suggest less clear-cut picture, with DA being involved also in performance directly (e.g. attention and WM) (Wang et al., 2004; Vijayraghavan et al., 2007; Shiner et al., 2012; Van Opstal et al., 2014). The goal of a simpler characterization was to elucidate how the two neuromodulators can influence each other for learning (DA) and performance (Ne). However, we stress that the ACC_Boost_ control mechanism could be easily, and without further assumptions, extended to DA modulation in the mesocortical dopaminergic pathway, for performance control in synergy with Ne (thus accounting for the more complex picture given by available empirical evidence).

The second limitation is the separation of the LC functions of learning rate modulation and cognitive control exertion. The cost of this separation between these two functions is outweighed by stable approximate optimal control of learning rate and catecholamines boosting policy. It must be stressed that the ACC_Boost_ module receives the LC signal *λ* related to learning rate in any case, making the boosting policy adaptive to environmental changes.

#### Relationship to other models and the central role of RL in dACC function

The RML belongs to a set of computational models suggesting RL as main function of mammalian dACC. Both the RVPM (RML direct predecessor; Silvetti et al., 2011) and the PRO model (Alexander and Brown, 2011) shares with the RML the main idea of the dACC as a state-action-outcome predictor. The RVPM is a subset of the dynamical version of the RML model (Model description: dynamical form, in Supplementary Methods). This implies that the RML can simulate also the results obtained by the RVPM (e.g., conflict monitoring, error likelihood estimation), extending even further the amount of empirical data that can be explained by this framework. Although the RML goes beyond these earlier works, by implementing meta-learning and higher-order conditioning, it shares with them the hypothesis that PE plays a core role for learning and decision-making. Indeed, we hypothesize that PE is a ubiquitous computational mechanism, which allows both dACC operations (equations 1 and 3) and the approximation of optimal learning rate in the LC (equations 6a-d).

Recent computational neuroscience of RL and decision-making focused on hierarchical architectures. For instance, Alexander and Brown (2015) proposed a hierarchical RL model (based on their previous PRO model), where hierarchical design is implemented within the dACC, unfolding in parallel with a hierarchical model of the DLPFC. In this model, PE afferents from hierarchically lower dACC layers work as an outcome proxy to train higher layers; at the same time, error predictions formulated at higher layers of DLPFC modulate outcome predictions at lower ones. This architecture successfully learned tasks where information is structured at different abstraction levels (like the 1-2AX task), exploring the RL basis of autonomous control of information access to WM.

In line with the recent interest for hierarchical RL, also Holroyd and McClure (2015) proposed a model exploiting hierarchical RL architecture (the HRL), where the dorsal striatum played a role of action selector, the dACC of task selector and the prelimbic cortex (in rodents) of context selector (where and when to execute a task). Moreover, each hierarchical layer implements a PE-based cognitive control signal that discounts option selection costs on the lower hierarchical level. This model can explain a wide variety of data about task selection and decision-making in cognitive and physical effort regulation.

The RML differs from these two models for the following reasons. i) Although higher-order conditioning in the RML is based, to some extent, on a hierarchical organization of reward signals (where DA signals linked to lower-level conditioned stimuli work as reward proxies to train higher-level stimuli), the RML lacks a genuine hierarchical structure. Its dynamics is emergent from the circular interaction between cortical and subcortical circuits, allowing meta-learning. ii) The RML provides mechanistic explanations to experimental findings from a broader range of domains (from effort modulation to higher-order conditioning), including findings that were earlier explained by the predecessor model RVPM (e.g. error processing, error likelihood estimation, etc.). iii) Both HER and HRL needed ad hoc parameter tuning to simulate different experimental results, while the RML (in both its dynamical and discrete versions) required just one single parameter set.

The RML represents cognitive control as dynamic selection of effort exertion, a mechanism that has been recently studied also by Verguts et al. (2015). In the latter model, effort exertion was dynamically optimized by the dACC as a process of RL-based decision-making, so that effort levels were selected to maximize long-term reward. This solution successfully simulated many experimental results from cognitive control and effort investment. A second model by Verguts (2017) described how dACC could implement cognitive control by functionally binding two or more brain areas by bursts of theta waves, whose amplitude would be proportional to the level of control. This theory describes how but not when (and neither how much) control should be exerted. The mechanisms proposed in the RML are an excellent complement to this theory, hypothesizing how, when and to what extent the dACC itself can decide to modulate theta bursts amplitude. It is worth noting that, in a very recent work, the PRO model provided a pure RL interpretation of dACC activation in preparation for an effortful task (Vassena et al., 2017a), where dACC activation preceding effortful action is due to effort intensity prediction and not to value of exerting effort or to any effort-related control signal. Although this theory is notable for parsimony, it does not provide any explanation about autonomous control of effort exertion.

Finally, Khamassi et al. (2011) also hypothesizes a role of dACC in meta-learning. The authors proposed a neural model (embodied in a humanoid robotic platform) where the temperature of the action selection process (i.e. the parameter controlling the trade-off between exploration and exploitation) was dynamically regulated as a function of PE signals. Like in the RML, dACC plays both a role in reward-based decision-making and in autonomous control of parameters involved in decision-making itself. Differently from the RML, this model provided a more classical view on PE origin, which were generated by the VTA and not by the dACC like in the RML. Moreover, mechanism proposed for temperature control was reactive to overall environmental variance (PE), lacking the capability to disentangle noise from volatility.

Concerning control of learning rate, earlier *Bayesian models* (e.g. Kalman 1960; Behrens et al. 2007; Mathys et al. 2011) also adapted their learning rates, proposing a computational account of behavioural adaptation. The main limitations of those models are their loose anatomical-functional characterization, the fact that they are computationally hard (in particular for optimal Bayesian solutions, e.g. Behrens et al. 2007), and the need of hyper-parameters specifying a priori the statistical structure of the environment (Kalman, 1960; Mathys et al., 2011). The latter feature reflects the peculiarity of these models of needing an ad-hoc hierarchical forward model representing the environment. At the top of this hierarchy, the experimenter defines a-priori crucial characteristics about the environment (like the precision of the probability function describing environmental volatility). At the best of our knowledge, the only Bayesian model able to estimate volatility without the need of specifying hyper-parameters is the one by Behrens et al. (2007), which works only for binary outcomes. Also Wilson et al., (2013) proposed an approximate Bayesian estimator that is based on PE, like the RML, without the need of specifying a forward model of the environment. However, the authors provided a solution for one subclass of volatility estimation problems (the change-point problems) and also in this case, an a priori hyper-parameter describing volatility (the process variance) was needed. In contrast, the RML provides an explicit neurophysiological theory on how near-optimal control emerges from the dialogue between dACC and LC, without the need of an a priori forward model and of information about environmental volatility itself. Indeed, our hyper-parameter *α* represents the minimal assumption that noise variance occurs at higher frequencies than process variance (volatility). This derives from the fact that the RML is not built hierarchically, but instead different interlocking modules influence one another, creating a circular interaction. Moreover, the RML can adapt learning rate in any kind of problem (binary, continuous, etc.), and finally, it integrates approximate Bayesian optimization with other cognitive functions, like effort control and higher-order conditioning.

#### Experimental predictions

The flexibility of RML, and the explicit neurophysiological hypotheses on which it is based, allow several experimental predictions. Here we list some potential experiments deriving from RML predictions. The first three could potentially falsify the model.

First, the RML architecture suggests that PE signal are generated by the dACC and then they converge toward the brainstem nuclei. This hypothesis implies two experimental predictions following dACC lesion: a) disruption of DA dynamics in higher-order conditioning, coupled with impossibility of higher-order instrumental training; b) disruption of LC dynamics related to learning rate control, coupled with behavioural flexibility impairment.

Second, a neurophysiological prediction concerns the mechanisms subtending higher-order conditioning and the difference between classical and instrumental paradigms. In the RML, higher-order conditioning is possible only when the agent plays an active role in learning (i.e., instrumental conditioning). We predict that hijacking the dACC decision of boosting catecholamines (e.g., via optogenetic intervention) would make possible higher-order conditioning in classical paradigms (ref. simulations 3a-b).

Third, in Simulation 2a (Figure 5b), the DA-lesioned RML predicts stronger dACC activation during an easy task (without effort) in presence of a high reward. This mechanism has been interpreted as a compensatory phenomenon allowing to avoid apathy (i.e. refusal to engage the task) if a small effort can make available a big reward. This is an explicit experimental prediction that could be tested both in animal paradigms and in mesolimbic DA impaired humans.

Furthermore, the model provides a promising platform for investigating the pathogenesis of several psychiatric disorders. In a previous computational work, we proposed how motivational and decision-making problems in attention-deficit/hyperactivity disorder (ADHD) could originate from disrupted DA signals to the dACC (Silvetti et al., 2013c). In the current paper, we also simulated a deficit related to cognitive effort (Simulation 2c) in case of DA deficit. Together, these findings suggest how DA deficit can cause both motivational and cognitive impairment in ADHD, with an explicit prediction on how DA deficit can impair also Ne dynamics (Hauser et al., 2016) in ADHD. This prediction could be tested by measuring performance and LC activation during decision-making or working memory tasks, while specifically modulating DA transmission in both patients (via pharmacological manipulation) and RML.

Another result with potential translational implication comes from Simulation 2 (and 2b in Supplementary Results), where the model suggested a possible mechanism linking boosting disruption and catecholamines dysregulation. This could be suggestive of pathogenesis of some depressive symptoms. More specifically, the RML predicts that DA antagonization intensifies effort in easy tasks (making them de facto subjectively harder) and decreases it in harder tasks (simulating apathy when effort is required by the environment; Figure 3b). Furthermore, it predicts an increased probability to refuse executing the task (thus simulating apathy). This effect could be experimentally tested by comparing effort-related dACC activation and behavioral patterns in tasks implying high and low effort with or without DA impairment. Another clinical application concerns a recent theory on autism spectrum disorder (ASD) pathogenesis. (Van de Cruys et al., 2014) proposed that a substantial number of ASD symptoms could be explained by dysfunctional control of learning rate and chronically elevate Ne release. This qualitative hypothesis could be easily implemented and explored quantitatively by altering learning rate meta-learning mechanisms in the RML leading to chronically high learning rate and LC activation.

#### Future perspectives

The RML shows how meta-learning involving three interconnected neuro-cognitive domains can account for the flexibility of the mammalian brain. However, our model is not meant to cover all aspects of meta-learning. Many other parameters may be meta-learned too. One obvious candidate is the temperature parameter of the softmax decision process (Khamassi et al., 2015), which arbitrates the exploration/exploitation trade-off. We recently proposed that this parameter is similarly meta-learned trading off effort costs versus rewards (Verguts et al., 2015). It must be noted that experimental findings indicated a link between LC activation and the arbitration on exploration/exploitation trade-off (Jepma and Nieuwenhuis, 2011), suggesting that the same mechanism used for earning rate optimization could be extended also to this domain. Other parameters from the classical RL modeling include discounting rate or eligibility traces (Schweighofer and Doya, 2003); future work should investigate the computational and biological underpinnings of their optimization.

Given the exceptionally extended dACC connectivity (Devinsky et al., 1995), other brain areas are likely relevant for the implementation of decision making in more complex settings. For example, we only considered model-free dynamics in RL and decision-making. However, both humans and nonhuman animals can rely also on complex environment models to improve learning and decision making (e.g. spatial maps for navigation or declarative rules about environment features). In this respect, future work should particularly focus on dACC-DLPFC-hippocampus interactions (Womelsdorf et al., 2014; Stoll et al., 2016), in order to investigate how environment models can modulate reward expectations and how the nervous system can represent and learn decision tree navigation (Pfeiffer and Foster, 2013).

Finally, the dynamical version of the RML functions in continuous time and in the presence of noise. These features are crucial to make a model survive outside the simplified environment of trial-level simulations, and make possible to simulate behaviour in the real world, like, for example, in robotic platforms (in preparation). RML embodiment into robotic platforms could be useful for both neuroscience and robotics. Indeed, testing our model outside the simplified environment of computer simulations could reveal model weaknesses that are otherwise hidden. Moreover, closing the loop between decision-making, body and environment (Pezzulo et al., 2011) is important to have a complete theory on the biological and computational basis of decision-making in the mammalian brain. At the same time, the RML could suggest new perspectives on natural-like flexibility in machine learning, helping, for example, in optimizing plasticity as a function of environmental changes.

Summing up, we formulated a model of how dACC-midbrain interactions may implement meta-learning in a broad variety of tasks. Besides understanding extant data and providing novel predictions, it holds the promise of taking cognitive control and, more in general, adaptive behaviour out of the experimental psychology lab and into the real world.

## Acknowledgements

Conflicts of interest: None. Thanks are due to Clay Holroyd, Gianluca Baldassarre, Daniele Caligiore, Giovanni Pezzulo, and Domenico Maisto for useful comments on this project. EV was supported by H2020 Marie Sklodowska-Curie Actions, project PreMotive, number 705630. EA was supported by Research Foundation Flanders under contract number 12C4715N.

- Bayesian flexibility of control

Lesione LC distrugge control LR

## Reference

Alexander WH, Brown JW (2011) Medial prefrontal cortex as an action-outcome predictor. Nat Neurosci 14:1338–1344.

Alexander WH, Brown JW (2015) Hierarchical Error Representation: A Computational Model of Anterior Cingulate and Dorsolateral Prefrontal Cortex. Neural Comput 27:2354–2410.

Apps MAJ, Ramnani N (2014) The Anterior Cingulate Gyrus Signals the Net Value of Others’ Rewards. J Neurosci 34:6190–6200.

Ashby FG, Ell SW, Valentin V V., Casale MB (2005) FROST: A Distributed Neurocomputational Model of Working Memory Maintenance. J Cogn Neurosci 17:1728–1743.

Aston-Jones G, Cohen JD (2005) An integrative theory of locus coeruleus-norepinephrine function: adaptive gain and optimal performance. Annu Rev Neurosci 28:403–450.

Behrens TE, Woolrich MW, Walton ME, Rushworth MF (2007) Learning the value of information in an uncertain world. Nat Neurosci 10:1214–1221.

Borst JP, Anderson JR (2013) Using model-based functional MRI to locate working memory updates and declarative memory retrievals in the fronto-parietal network. Proc Natl Acad Sci 110:1628–1633.

Chong TT-J, Apps M, Giehl K, Sillence A, Grima LL, Husain M (2017) Neurocomputational mechanisms underlying subjective valuation of effort costs Seymour B, ed. PLOS Biol 15:e1002598.

D’Esposito M, Postle BR (2015) The Cognitive Neuroscience of Working Memory. Annu Rev Psychol 66:115–142.

Denny M, Ratner S (1970) Comparative psychology: Research in animal behavior. Oxford: Dorsey Press.

Devinsky O, Morrell MJ, Vogt BA> (1995) Contributions of anterior cingulate cortex to behaviour. Brain 118 (Pt 1:279–306.

Doya K (2002) Metalearning and neuromodulation. Neural Netw 15:495–506.

Ebitz RB, Hayden BY (2016) Dorsal anterior cingulate: a Rorschach test for cognitive neuroscience. Nat Neurosci 19:1278–1279.

Frank MJ, Seeberger LC, O’Reilly R C (2004) By carrot or by stick: cognitive 45 reinforcement learning in parkinsonism. Science (80-) 306:1940–1943.

Grossberg S (1980) How does a brain build a cognitive code? Psychol Rev 87:1–51.

Hauser TU, Fiore VG, Moutoussis M, Dolan RJ (2016) Computational Psychiatry of ADHD: Neural Gain Impairments across Marrian Levels of Analysis. Trends Neurosci 39:63–73.

Holroyd CB, Coles MG (2002) The neural basis of human error processing: reinforcement learning, dopamine, and the error-related negativity. Psychol Rev 109:679–709.

Holroyd CB, McClure SM (2015) Hierarchical control over effortful behavior by rodent medial frontal cortex: A computational model. Psychol Rev 122:54–83.

Jepma M, Murphy PR, Nassar MR, Rangel-Gomez M, Meeter M, Nieuwenhuis S (2016) Catecholaminergic Regulation of Learning Rate in a Dynamic Environment. O’Reilly JX, ed. PLoS Comput Biol 12:e1005171.

Jepma M, Nieuwenhuis S (2011) Pupil Diameter Predicts Changes in the Exploration-Exploitation Tradeoff: Evidence for the Adaptive Gain Theory. J Cogn Neurosci 23:1587–1596.

Joshi S, Li Y, Kalwani RM, Gold JI (2016) Relationships between Pupil Diameter and Neuronal Activity in the Locus Coeruleus, Colliculi, and Cingulate Cortex. Neuron 89:221–234.

Kahneman D (1973) Attention and effort. Prentice-Hall.

Kalman R (1960) A new approach to linear filtering and prediction problems. J basic Eng.

Kennerley SW, Behrens TE, Wallis JD (2011) Double dissociation of value computations in orbitofrontal and anterior cingulate neurons. Nat Neurosci 14:1581–1589.

Khamassi M, Lallée S, Enel P, Procyk E, Dominey PF (2011) Robot cognitive control with a neurophysiologically inspired reinforcement learning model. Front Neurorobot 5:1.

Khamassi M, Quilodran R, Enel P, Dominey PF, Procyk E (2015) Behavioral Regulation and the Modulation of Information Coding in the Lateral Prefrontal and Cingulate Cortex. Cereb Cortex 25:3197–3218.

Kolling N, Wittmann MK, Behrens TEJ, Boorman ED, Mars RB, Rushworth MFS (2016) Value, search, persistence and model updating in anterior cingulate cortex. Nat Neurosci 19:1280–1285.

Kool W, Botvinick M (2013) The intrinsic cost of cognitive control. Behav Brain Sci 36:661–698.

Kool W, McGuire JT, Rosen ZB, Botvinick MM (2010) Decision making and the avoidance of cognitive demand. J Exp Psychol Gen 139:665–682.

Langner R, Eickhoff SB (2013) Sustaining attention to simple tasks: a meta-analytic review of the neural mechanisms of vigilant attention. Psychol Bull 139:870–900.

Li BM, Mao ZM, Wang M, Mei ZT (1999) Alpha-2 adrenergic modulation of prefrontal cortical neuronal activity related to spatial working memory in monkeys. Neuropsychopharmacology 21:601–610.

Li BM, Mei ZT (1994) Delayed-response deficit induced by local injection of the alpha 2-adrenergic antagonist yohimbine into the dorsolateral prefrontal cortex in young adult monkeys. Behav Neural Biol 62:134–139.

Ljungberg T, Apicella P, Schultz W (1992) Responses of monkey dopamine neurons during learning of behavioral reactions. J Neurophysiol 67:145–163.

Margulies DS, Kelly AMC, Uddin LQ, Biswal BB, Castellanos FX, Milham MP (2007) Mapping the functional connectivity of anterior cingulate cortex. Neuroimage 37:579–588.

Mathys C, Daunizeau J, Friston KJ, Stephan KE (2011) A Bayesian foundation for individual learning under uncertainty. Front Hum Neurosci 5:39.

Nassar MR, Rumsey KM, Wilson RC, Parikh K, Heasly B, Gold JI (2012) Rational regulation of learning dynamics by pupil-linked arousal systems. Nat Neurosci 15:1040–1046.

Niv Y, Daw ND, Joel D, Dayan P (2007) Tonic dopamine: opportunity costs and the control of response vigor. Psychopharmacology (Berl) 191:507–520.

Niv Y, Joel D, Dayan P (2006) A normative perspective on motivation. Trends Cogn Sci 10:375–381.

O’Reilly RC, Frank MJ, Hazy TE, Watz B (2007) PVLV: the primary value and learned value Pavlovian learning algorithm. Behav Neurosci 121:31–49.

Pezzulo G, Barsalou L, Cangelosi A (2011) The mechanics of embodiment: A dialog on embodiment and computational modeling. Embodied and.

Pezzulo G, Rigoli F, Chersi F (2013) The Mixed Instrumental Controller: Using Value of Information to Combine Habitual Choice and Mental Simulation. Front Psychol 4:92.

Pfeiffer BE, Foster DJ (2013) Hippocampal place-cell sequences depict future paths to remembered goals. Nature 497:74–79.

Pierce W, Cheney D (2004) Behavior Analysis and Learning New Jersey: Laurence Erlbaum Associates.

Rushworth MF, Behrens TE (2008) Choice, uncertainty and value in prefrontal and cingulate cortex. Nat Neurosci 11:389–397.

Salamone JD, Cousins MS, Bucher S (1994) Anhedonia or anergia? Effects of haloperidol and nucleus accumbens dopamine depletion on instrumental response selection in a T-maze cost/benefit procedure. Behav Brain Res 65:221–229.

Sara SJ (2009) The locus coeruleus and noradrenergic modulation of cognition. Nat Rev Neurosci 10:211–223.

Schultz W, Apicella P, Ljungberg T (1993) Responses of monkey dopamine neurons to reward and conditioned stimuli during successive steps of learning a delayed response task. J Neurosci 13:900–913.

Schweighofer N, Doya K (2003) Meta-learning in reinforcement learning. Neural Netw 16:5–9.

Shenhav A, Botvinick MM, Cohen JD (2013) The expected value of control: an integrative theory of anterior cingulate cortex function. Neuron 79:217–240.

Shenhav A, Cohen JD, Botvinick MM (2016) Dorsal anterior cingulate cortex and the value of control. Nat Neurosci 19:1286–1291.

Shenhav A, Straccia MA, Cohen JD, Botvinick MM (2014) Anterior cingulate engagement in a foraging context reflects choice difficulty, not foraging value. Nat Neurosci 17:1249–1254.

Shiner T, Seymour B, Wunderlich K, Hill C, Bhatia KP, Dayan P, Dolan RJ (2012) Dopamine and performance in a reinforcement learning task: evidence from Parkinson’s disease. Brain 135:1871–1883.

Silvetti M, Alexander W, Verguts T, Brown JW (2014) From conflict management to reward-based decision making: Actors and critics in primate medial frontal cortex. Neurosci Biobehav Rev 46:44–57.

Silvetti M, Seurinck R, van Bochove ME, Verguts T (2013a) The influence of the noradrenergic system on optimal control of neural plasticity. Front Behav Neurosci in press:160.

Silvetti M, Seurinck R, Verguts T (2011) Value and prediction error in medial frontal 48 cortex: integrating the single-unit and systems levels of analysis. Front Hum Neurosci 5:75.

Silvetti M, Seurinck R, Verguts T (2013b) Value and prediction error estimation account for volatility effects in ACC: A model-based fMRI study. Cortex.

Silvetti M, Wiersema JR, Sonuga-Barke E, Verguts T (2013c) Deficient reinforcement learning in medial frontal cortex as a model of dopamine-related motivational deficits in ADHD. Neural Netw 46:199–209.

Stoll FM, Fontanier V, Procyk E (2016) Specific frontal neural dynamics contribute to decisions to check. Nat Commun 7:11990.

Van de Cruys S, Evers K, Van der Hallen R, Van Eylen L, Boets B, de-Wit L, Wagemans J (2014) Precise minds in uncertain worlds: Predictive coding in autism. Psychol Rev 121:649–675.

Van Opstal F, Van Laeken N, Verguts T, van Dijck J-P, De Vos F, Goethals I, Fias W (2014) Correlation between individual differences in striatal dopamine and in visual consciousness. Curr Biol 24:R265–R266.

Varazzani C, San-Galli A, Gilardeau S, Bouret S (2015) Noradrenaline and dopamine neurons in the reward/effort trade-off: a direct electrophysiological comparison in behaving monkeys. J Neurosci 35:7866–7877.

Vassena E, Deraeve J, Alexander W (2017a) Predicting motivation: computational models of PFC can explain neural coding of motivation and effort-based decision-making in health and disease. J Cogn.

Vassena E, Holroyd C, Alexander WH (2017b) Computational models of anterior cingulate cortex: At the crossroads between prediction and effort. Front Neurosci 11.

Vassena E, Silvetti M, Boehler CN, Achten E, Fias W, Verguts T (2014) Overlapping Neural Systems Represent Cognitive Effort and Reward Anticipation Maurits NM, ed. PLoS One 9:e91008.

Verguts T (2017) Binding by Random Bursts: A Computational Model of Cognitive Control. J Cogn Neurosci 29:1103–1118.

Verguts T, Vassena E, Silvetti M (2015) Adaptive effort investment in cognitive and physical tasks: a neurocomputational model. Front Behav Neurosci 9:57.

Vijayraghavan S, Wang M, Birnbaum SG, Williams G V, Arnsten AF (2007) Inverted-U dopamine D1 receptor actions on prefrontal neurons engaged in working memory. Nat Neurosci 10:376–384.

Walsh MM, Anderson JR (2014) Navigating complex decision spaces: Problems and paradigms in sequential choice. Psychol Bull 140:466–486.

Walton ME, Groves J, Jennings KA, Croxson PL, Sharp T, Rushworth MFS, Bannerman DM (2009) Comparing the role of the anterior cingulate cortex and 6-hydroxydopamine nucleus accumbens lesions on operant effort-based decision making. Eur J Neurosci 29:1678–1691.

Wang M, Ramos BP, Paspalas CD, Shu Y, Simen A, Duque A, Vijayraghavan S, Brennan A, Dudley A, Nou E, Mazer JA, McCormick DA, Arnsten AFT (2007) α2A-Adrenoceptors Strengthen Working Memory Networks by Inhibiting cAMP-HCN Channel Signaling in Prefrontal Cortex. Cell 129:397–410.

Wang M, Vijayraghavan S, Goldman-Rakic PS (2004) Selective D2 receptor actions on the functional circuitry of working memory. Science (80-) 303:853–856.

Welch G, Bishop G (1995) An introduction to the Kalman filter.

Williams J, Dayan P (2005) Dopamine, learning, and impulsivity: a biological account of attention-deficit/hyperactivity disorder. J Child Adolesc Psychopharmacol 15:160–169.

Wilson RC, Nassar MR, Gold JI (2013) A Mixture of Delta-Rules Approximation to Bayesian Inference in Change-Point Problems Behrens T, ed. PLoS Comput Biol 9:e1003150.

Womelsdorf T, Ardid S, Everling S, Valiante TA (2014) Burst firing synchronizes prefrontal and anterior cingulate cortex during attentional control. Curr Biol 24:2613–2621.

Yu AJ (2007) Adaptive Behavior: Humans Act as Bayesian Learners. Curr Biol 17:R977–R980.

Yu AJ, Dayan P (2005) Uncertainty, neuromodulation, and attention. Neuron 46:681–692.

